# *Toxoplasma* induces stripping of perisomatic inhibitory synapses

**DOI:** 10.1101/788190

**Authors:** Gabriela L. Carrillo, Valerie A. Ballard, Taylor Glausen, Zack Boone, Joseph Teamer, Cyrus L. Hinkson, Elizabeth A. Wohlfert, Ira J. Blader, Michael A. Fox

## Abstract

Infection and inflammation within the brain induces changes in neuronal connectivity and function. The intracellular protozoan parasite, *Toxoplasma gondii*, is one pathogen that infects the brain and can cause encephalitis and seizures. Persistent infection by this parasite is also associated with behavioral alterations and an increased risk for developing psychiatric illness, including schizophrenia. Current evidence from studies in humans and mouse models suggest that both seizures and schizophrenia result from a loss or dysfunction of inhibitory synapses. In line with this, we recently reported that persistent *Toxoplasma gondii* infection alters the distribution of glutamic acid decarboxylase 67 (GAD67), an enzyme that catalyzes GABA synthesis in inhibitory synapses. These changes could reflect a redistribution of presynaptic machinery in inhibitory neurons or a loss of inhibitory nerve terminals. To directly assess the latter possibility, we employed serial block face scanning electron microscopy (SBFSEM) and quantified inhibitory perisomatic synapses in neocortex and hippocampus following parasitic infection. Not only did persistent infection lead to a significant loss of perisomatic synapses, it induced the ensheathment of neuronal somata by phagocytic cells. Immunohistochemical, genetic, and ultrastructural analyses revealed that these phagocytic cells included reactive microglia. Finally, ultrastructural analysis identified phagocytic cells enveloping perisomatic nerve terminals, suggesting they may participate in synaptic stripping. Thus, these results suggest that microglia contribute to perisomatic inhibitory synapse loss following parasitic infection and offer a novel mechanism as to how persistent *Toxoplasma gondii* infection may contribute to both seizures and psychiatric illness.

**MAIN POINTS:**

- *Toxoplasma*-infection leads the loss of perisomatic inhibitory synapses
- Phagocytic microglia ensheath neuronal somata following *Toxoplasma*-infection
- Microglia contact and envelop perisomatic nerve terminals, suggesting that *Toxoplasma* induces synaptic stripping

## INTRODUCTION

*Toxoplasma gondii* is a widespread obligate, intracellular protozoan parasite that infects roughly one-third of the world’s population (Pappas et al. 2009; Montoya and Liesenfeld 2004). The parasite reproduces sexually in the felid intestine or asexually in most warm-blooded animals. Following digestion of the resulting oocysts (generated by sexual reproduction) or tissue cysts (generated in intermediate host tissues), intestinal enzymes degrade wall of these structures freeing sporozoites or bradyzoites, respectively, which then infect intestinal epithelial cells and differentiate into virulent tachyzoites (Montoya and Liesenfeld 2004). Inflammatory cells recruited to the site of initial infection become infected by these tachyzoites and assist in their dissemination throughout the body of the intermediate host, including to the brain, retina and skeletal muscle (Lambert et al. 2006; Courret et al. 2006, Coombes et al. 2013). Within these tissues, tachyzoites convert to slowly dividing bradyzoites that develop into tissue cysts that may persist for the lifetime of the intermediate host.

In the brain, after the parasite triggers an initial inflammatory response, bradyzoite-containing cysts preferentially form within neurons (Sims et al. 1989; McConkey et al. 2013; Cabral et al. 2016). A growing body of literature suggests that persistent *Toxoplasma gondii* infection induces behavioral changes in intermediate hosts, including humans (Poirotte et al. 2016; Berdoy et al. 2000; Vyas et al. 2007; Beste et al. 2014; Stock et al. 2014). Furthermore, over four dozen studies have identified *Toxoplasma gondii* infection as a risk factor for developing schizophrenia (Wang et al. 2019, Xiao et al. 2018, Burgdorf et al. 2019; Dickerson et al. 2017, 2014, 2018; Kano et al. 2018), a complex disorder characterized by altered cognitive performance, the acquisition and expression of behaviors absent from healthy individuals (such as hallucinations), and the loss of behaviors normally present in healthy individuals (leading to social withdrawal, apathy, and neglect). A number of cellular and molecular mechanisms have been proposed to underlie behavioral changes in schizophrenia, many of which point to alterations in the assembly, maintenance and function of synapses (Birnbaum and Weinberger 2017). In addition to its association with schizophrenia, persistent *Toxoplasma* infection in animal models leads to seizures (Brooks et al. 2015; David et al. 2016), altered neurotransmitter levels (Alsaady et al. 2019; Skallova et al. 2006; Martin et al. 2015; Gatkowska et al. 2013), altered neural connectivity (Brooks et al. 2015; Ihara et al. 2016; Parlog et al. 2015; Lang et al. 2018) and altered neuronal function (Haroon et al. 2012), all phenotypes that are associated with impaired synaptic organization or function.

Synapses can be broadly categorized into those whose activity increases the probability of activity in a postsynaptic neuron (i.e. excitatory synapses) and those whose activity reduces the probability of activity in a postsynaptic neuron (i.e. inhibitory synapses). Although inhibitory synapses comprise only a fraction (~20%) of all synapses in neocortex or hippocampus, they are essential for controlling the flow of information transfer in the brain and their dysfunction has been associated with neurological and neuropsychiatric disorders (Marin 2012). In fact, we previously identified alterations in the distribution of glutamic acid decarboxylase 67 (GAD67), the enzyme required to generate the inhibitory neurotransmitter GABA, in *Toxoplasma*-infected mice (Brooks et al. 2015). These changes could reflect a redistribution of presynaptic machinery in inhibitory neurons or a loss of inhibitory nerve terminals. This motivated us to directly test whether *Toxoplasma gondii* infection leads to the loss of inhibitory synapses. We focused our attention on perisomatic inhibitory synapses formed by Parvalbumin-expressing, fast-spiking interneurons in the cerebral cortex and hippocampus as the loss or impairment of these specific synapses has been associated with seizures, schizophrenia, and schizophrenia-related behaviors (Schwaller et al. 2004; Belforte et al. 2010; Gonzalez-Burgos et al. 2010, 2011; Gonzalez-Burgos and Lewis 2012; Lewis et al. 2011; Wohr et al. 2015; Hamm et al. 2017; Mukherjee et al. 2019). Using serial block face scanning electron microscopy (SBFSEM) and immunolabeling, we discovered *Toxoplasma* infection leads to a significant loss of these perisomatic inhibitory synapses. Moreover, we observed ensheathment of neuronal somata (and perisomatic nerve terminals) by phagocytic microglia, suggesting that these cells may contribute to the loss of inhibitory synapses following long-term infection. Thus, these data suggest that *Toxoplasma* may increase the risk of developing seizures and schizophrenia by contributing to microglial activation and the loss of perisomatic inhibitory synapses.

## MATERIALS AND METHODS

### Animals

C57BL/6J mice were purchased from Jackson Laboratories (Bar Harbor, ME) and *Cx3cr1-GFP* mice (Jackson Laboratories, Stock# 005582) were provided by Dr. M. Theus (Virginia Tech). All analyses conformed to National Institutes of Health guidelines and protocols approved by the University at Buffalo and Virginia Polytechnic Institute and State University Institutional Animal Care and Use Committees.

### Parasite infections

Mice (8-12 weeks of age) were infected with type II strains of *Toxoplasma gondii* as previously described (Brooks et al. 2015). Brains from chronically infected mice (30 to 90 days post-infection) were isolated and homogenized in phosphate-buffered saline pH 7.4 (PBS). The total number of *Toxoplasma gondii* type II ME49 or RFP-expressing ME49 cysts within 20µl of brain homogenate was counted by brightfield or fluorescence microscopy, respectively. Mice were interperitoneally infected with 5 cysts of either in PBS. After 30 days, brains from *Toxoplasma gondii*-infected or mock-infected (which received IP injection of PBS) were collected for immunohistochemistry (IHC) or serial block face scanning electron microscopy (SBFSEM) analysis.

### Immunohistochemistry

IHC was performed as described previously (Su et al. 2010, 2016; Brooks et al. 2015). Anesthetized mice were transcardially perfused with PBS pH 7.4 followed by 4% paraformaldehyde in PBS (PFA; pH 7.4). Post-fixed brains (12-16 hrs at 4°C in 4% PFA) were harvested, cryopreserved in 30% sucrose for 3 days, and embedded in Tissue Freezing Medium (Electron Microscopy Sciences, Hatfield, PA). Brains were coronally cryostectioned at 16µm and were air dried for 15 min at room temperature before incubation with blocking buffer (5% normal goat serum-2.5% bovine serum albumin-0.1% Triton X-100-PBS) for 60 min. Primary antibodies were diluted in blocking buffer as follows: GAD67 1:500 (Millipore MAB5406); IBA1 1:800 (Wako, Richmond, VA; 019-19741); CD68 1:1000 (Abcam, Cambridge, UK; AB53444); NeuroTrace 1:300 in PBS (ThermoFisher, Waltham, MA; N-21480) and incubated on coronal brain sections overnight at 4°C. Slides were washed in PBS and incubated with fluorescently conjugated secondary antibodies obtained from Molecular Probes/Invitrogen (diluted 1:1,000 in blocking buffer). After washing in PBS, slides were mounted with Vectashield (Vector Laboratories, Burlingame, CA) and images were acquired on a Zeiss LSM 700 confocal microscope. Identical imaging parameters were used for the acquisition of images for different experimental conditions. A minimum of three animals and three image fields per brain region (per experimental condition) were compared in all experiments.

### Image analysis

For binary quantification of fluorescent images, single channels of each image obtained after immunohistochemistry (as described above) were converted to binary images and the percent field occupied by immunoreactivity was obtained using ImageJ software (see Brooks et al. 2015; Su et al. 2016; Monavarfeshani et al. 2018). For quantifying the ratio of immunoreactivity, the average fluorescent intensity was measured in images of stratum pyramidalis and in the surrounding hippocampal layers (SR and SO). DAPI labeling was used to differentiate these layers. Average fluorescent intensity was calculated using ImageJ software. Rations of the fluorescent intensities in these regions was calculated and a student’s t test was used to test for statistical significance. Quantification of the number of cells ensheathed by Iba1^+^ phagocytes was performed using maximum-projection images of merged channels. Ensheathed cells were identified by observing phagocytes (Iba1^+^ or GFP^+^ cells) that were both in direct contact with non-phagocytes (Iba1^−^/DAPI^+^ or GFP^−^/DAPI^+^ cells) and extended processes around a fraction of those cell somas. Examples of Iba1^+^ cells in contact with Iba1^−^/DAPI^+^ cells are shown by the arrows in Figure 5B,D and examples of GFP^+^ cells in contact with GFP^−^/DAPI^+^ cells are shown by the arrows in Figure 5F,L. Total ensheathed cells were counted in each field of view for each condition. For all analysis, six mice were used per condition with a minimum of three images per brain region.

### *In situ* hybridization (ISH)

ISH was performed on 16 µm coronal sections as described previously in (Su et al. 2010; Monavarfeshani et al. 2018). Briefly, tissue slides were incubated with diluted and heat-denatured *Gad1* and *Syt1* riboprobes (1-2µm of riboprobe in 60 µl of hybridization solution heated for 12 min in 70°C) overnight in 60°C incubation. Bound riboprobes were detected by staining with Tyramide Signal Amplification (TSA) system. Immunohistochemistry was immediately performed following TSA system signal detection as described above. Images were obtained on a Zeiss LSM700 confocal microscope. A minimum of three animals per condition were compared in ISH experiments.

### Serial block face scanning electron microscopy

SBFSEM was performed as described previously (Hammer et al. 2014; Monavarfeshani et al. 2018). Briefly, mice were perfused with 0.1M sodium cacodylate buffer containing 4% PFA and 2.5% glutaraldehyde and immediately 300 µm vibratomed coronal sections were collected. CA1 regions of hippocampus and layer V of neocortex were microdissected and sent to Renovo Neural (Cleveland, OH) for staining, processing and image acquisition. Datasets represented regions that were 50µm x 50-150µm and consisted of ~500 serial sections (with each section being 50-60 nm thick). SBFSEM images were imaged at a 5nm/pixel resolution. Serial image stacks were manually traced and analyzed using TrakEM2 in Fiji (Cardona et al., 2012).

## RESULTS

### Toxoplasma infection leads to the loss of perisomatic inhibitory synapses

To assess how parasitic infection affects GABAergic inhibitory synapses in the mammalian brain, adult mice were mock-infected or infected with the type II ME49 *Toxoplasma* strain. Thirty days after infection, brains were harvested and GABAergic synapses in hippocampus and neocortex were analyzed. These regions were selected since they contain a high concentration of perisomatic inhibitory synapses formed by fast-spiking, Parvalbumin (Parv)-expressing interneurons. In mock-infected controls, punctate GAD67-immunoreactivity labels inhibitory synapses throughout the brain, however, in the Cornu Ammonis (CA) regions of hippocampus, the majority of these inhibitory synapses are concentrated in stratum pyramidalis (SP) (Figure 1A,B). In neocortex, GAD67-labeled inhibitory synapses are distributed in all cortical layers, although an increased density is observed in layer V (and in layer II/III), where a high proportion of these synapses surround neuronal cell bodies (Figure 1E,F). Similar to what we previously observed in the CA3 subregion of ME49-infected hippocampus (Brooks et al. 2015), data presented revealed GAD67 immunoreactivity is less punctate and more diffusely localized in other hippocampal regions and in neocortex following ME49 infection (Figure 1A-K, and Supplemental Figure 1). Not only did we observe this redistribution of GAD67 immunoreactivity, we observed a loss of GAD67^+^ terminals surrounding neuronal somas in ME49-infected tissues (Figure 1L-O). We interpret this data to suggest a loss of GABAergic perisomatic synapses following persistent *Toxoplasma* infection.

**Figure 1.**
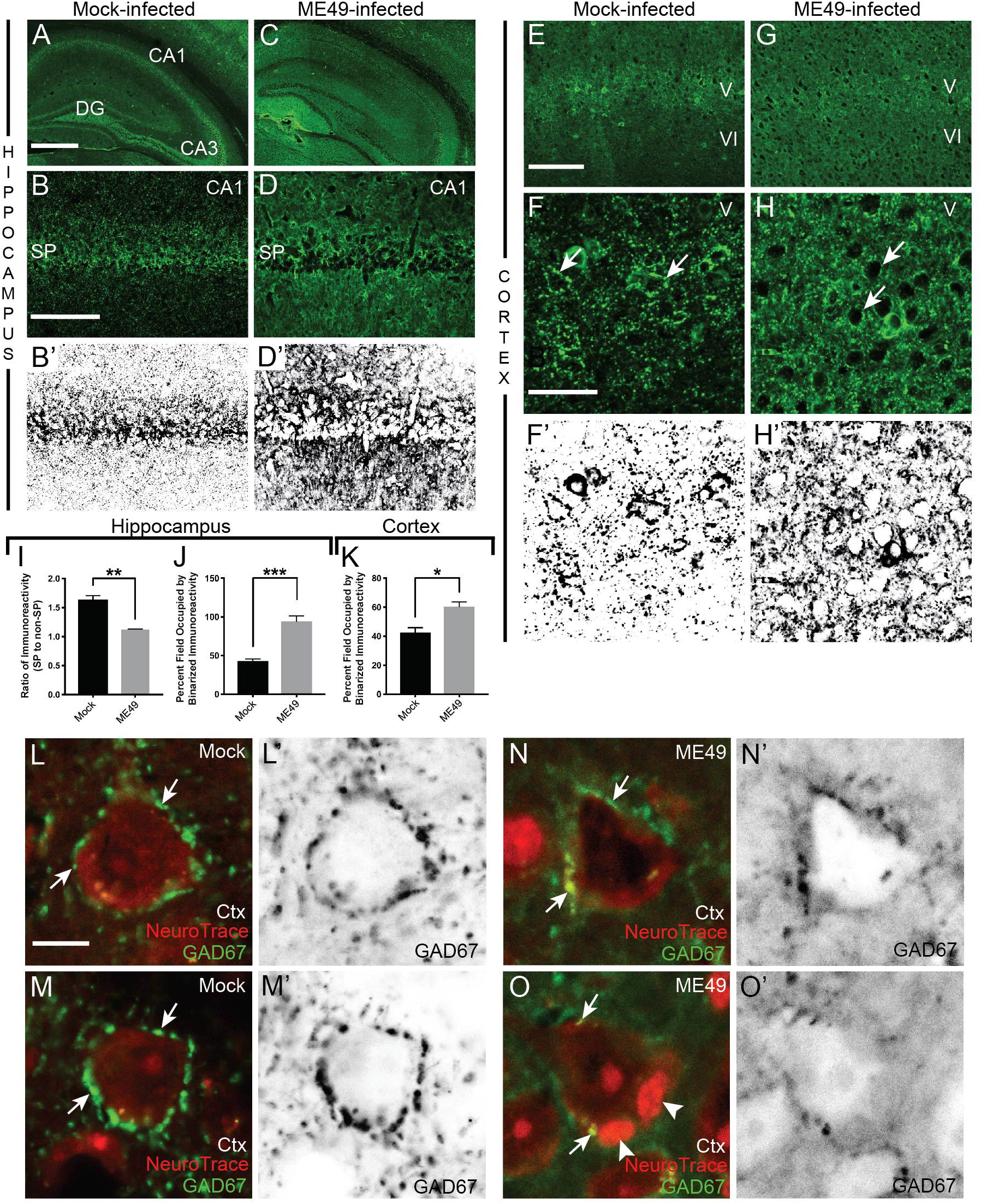
Disruption of inhibitory synapses in hippocampus and neocortex of *Toxoplasma*-infected mice. **A-D.** GAD67-immunostaining of inhibitory synapses in mock-(**A**) and ME49-(**C**) infected hippocampi. Hippocampal regions CA1, CA3, and dentate gyrus (DG) are shown in **A**. ME49 infection induced a redistribution of GAD67-immunoreactivity, as can be seen in high magnification images of the CA1 region of hippocampus (**B,D**). Binarized, inverted images of GAD67-immunoreactivity in CA1 shows a concentration of inhibitory synapses in stratum pyramidalis (SP) of mock-infected tissue (**B’**). In contrast, this signal is redistributed out of SP in ME49-infected mice (**D’**). **E-H**. GAD67-immunostaining of inhibitory synapses in layer V of mock-(**E**) and ME49-(**G**) infected neocortex. High magnification images of layer V in mock-infected tissue (**F**) shows GAD67^+^ perisomatic synapses clustering around cell bodies (arrows). Fewer GAD67^+^ perisomatic synapses are present in ME49-infected layer V (**H**). **F’,H’** show binarized, inverted images of GAD67-immunoreactivity in layer V of mock- and ME49-infected mice. **I**. Quantification of GAD67-immunoreactivity in SP (versus non-SP regions) of the CA1 region of hippocampus of mock and ME49-infected mice. Bars represent means +/− SEM. ** indicates p<0.01 by Students T-test (n=6). **J,K.** Quantification of the area of binarized, inverted GAD67-immunoreactivity in CA1 (**J**) and layer V of neocortex (**K**) of mock and ME49-infected mice. Bars represent means +/− SEM. * indicates p<0.05 and *** indicates p<0.001 by Students T-test (n=3). **L-O.** High magnification images of GAD67^+^ inhibitory terminals on NeuroTrace-labeled somas in layer V of mock-**(L,M)** and ME49-**(N,O)** neocortex. **I’-L’** show binarized, inverted images of GAD67-immunoreactivity in mock-**(L’,M’)** and ME49-**(N’,O’)** neocortex. Arrows indicate GAD67^+^ perisomatic synapses. Arrowheads in **L** highlight two small, NeuroTrace-labeled cells contacting the soma of a pyramidal neuron. Scale bar in **A** = 500 µm for **A,C**; in **B** = 100 µm for **B,B’,D,D’**; in **E** = 200 µm for **E,G**; in **F** = 50 µm for **F,F’,H,H’**; in **L** = 10 µm for **L-O**; **L’-O’**.

In both stratum pyramidalis of the hippocampus and layer V of cerebral cortex, Parv^+^ interneurons generate GAD67^+^ perisomatic synapses onto both excitatory pyramidal neurons and other Parv^+^ inhibitory interneurons (Supplemental Figure 1) (Pi et al. 2013; Pfeffer et al. 2013). To assess whether the loss of GAD67^+^ perisomatic synapses was cell-specific or occurred on both excitatory and inhibitory neurons, we immunolabeled GAD67^+^ terminals and performed *in situ* hybridization for either *Synaptotagmin1* (*Syt1*) mRNA or *Glutamic Acid Decarboxylase 1* (*Gad1)* mRNA (which encodes GAD67) to discriminate between excitatory and inhibitory postsynaptic neurons, respectively. *Syt1* mRNA is transcribed by a large subset of excitatory neurons (and some interneurons) in stratum pyramidalis of CA1 and layer V of cortex (Marqueze et al. 1995; Su et al. 2010). In contrast, *Gad1* mRNA is present in most, if not all, inhibitory neurons (and in no excitatory neurons) in these regions. We observed abundant GAD67^+^ nerve terminals surrounding both *Syt1*^*+*^ or *Gad1*^*+*^ neurons in controls (Supplemental Figure 1), but few GAD67^+^ terminals were observed surrounding either populations of neurons in *Toxoplasma*-infected mice (Supplemental Figure 1). These data suggest a loss of GABAergic perisomatic synapses onto both pyramidal neurons and interneurons in hippocampus and neocortex of *Toxoplasma*-infected brains.

Unfortunately, quantifying these observations proved challenging since it was unclear with this resolution of analysis as to whether GAD67^+^ terminals truly formed synaptic connections onto the adjacent somas. This was particularly true in the hippocampus where pyramidal neurons are densely packed in stratum pyramidalis. Moreover, it was unclear whether the absence of GAD67^+^ terminals reflected the loss of perisomatic synapses or merely the loss of GAD67-immunoreactivity within these nerve terminals. To circumvent these issues, we employed serial block face scanning electron microscopy (SBFSEM), which allows for large volumes of serial, ultrastructural data to be collected and analyzed in 3D (Denk and Horsfmath 2014). SBFSEM datasets were generated for both CA1 and neocortex of mock-infected and ME49-infected mice. At low magnification, we identified regions of stratum pyramidalis (SP) in CA1 or layer V in neocortex to analyze in detail (Figure 2A,B, and data not shown). Once regions of interest were selected, serial sections were obtained at high resolution (5nm/pixel) (Figure 2C). SBFSEM images resemble data from traditional transmission electron microscopy allowing us to use traditional ultrastructural morphological criteria to identify neurons, glial cells, blood vessels, synapses and parasite cysts (Figure 2C) (Savage et al. 2018; Chen et al. 2014).

**Figure 2.**
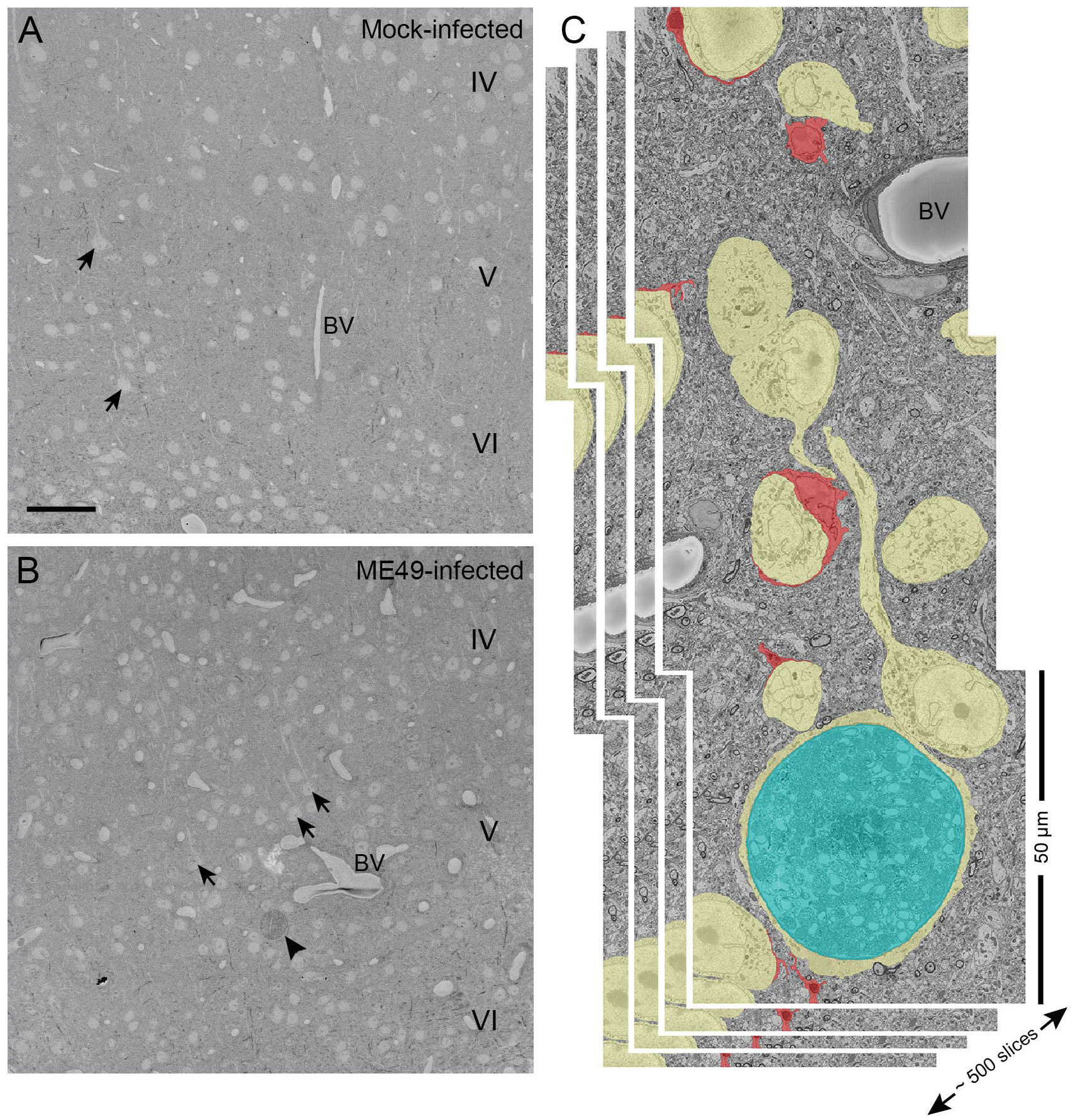
Serial block face scanning electron microscopic (SBFSEM) analysis of mock- and ME49-infected brain tissue. **A, B.** Low magnification SBFSEM images of cerebral cortex of mock-(**A**) and ME49-(**B**) infected mice. Cortical layers IV, V, and VI are labeled. Arrows highlight layer V pyramidal neurons. Arrowhead in **B** highlights a *Toxoplasma gondii* cyst. BV – blood vessels. Scale bar in **A** = 50 µm for **A,B**. **C.** High magnification montage of 3 adjacent SBFSEM datasets. Each dataset consists of ~500 serial 50 µm x 50 µm micrographs, each 60 nm thick. In these micrographs, neurons are pseudocolored yellow, microglia are red, and a *Toxoplasma gondii* cyst is pseudocolored blue.

Perisomatic synapses in hippocampus and cerebral cortex were identified by the presence of mitochondria, synaptic vesicles, and active zones that opposed adjacent neuronal somata (Figure 3A-H). To quantify perisomatic synapse number, we identified nine neurons in mock- and ME49-infected stratum pyramidalis whose entire somata were present in the 3D SBFSEM datasets and manually traced all of the perisomatic synapses onto these neurons (Figure 3A-D,I). Our analyses revealed that infection significantly reduced the numbers of hippocampal perisomatic synapses (Figure 3I). Persistent parasitic infection has been shown to cause neuronal changes, including neuronal loss due to excitotoxicity (David et al. 2016; Torres et al. 2018), therefore we also addressed the possibility that the decrease in perisomatic synapses was due to changes in neuronal size. We analyzed the number of serial sections per neuron in mock- and ME49-infected hippocampus and observed a modest decrease in neuronal size following infection (Figure 3J). However, even when we accounted for this difference, a significant reduction in the ratio of perisomatic synapses per section was observed in stratum pyramidalis (Figure 3K).

**Figure 3.**
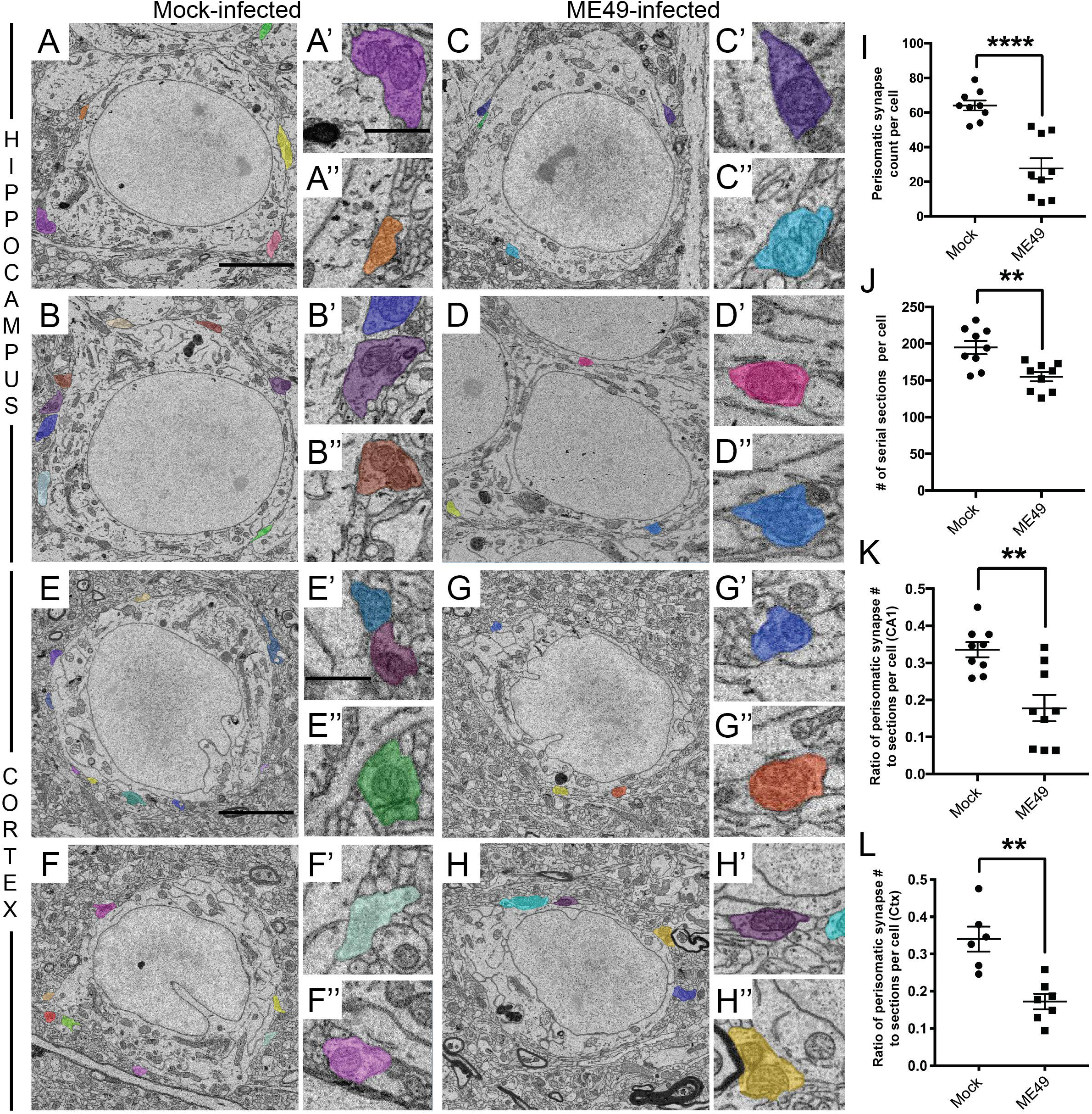
*Toxoplasma* infection leads to a loss of perisomatic synapses in hippocampus and layer V of neocortex. **A-D.** SBFSEM micrographs of neurons in stratum pyramidalis of the CA1 region of hippocampus of mock-(**A,B**) and ME49-(**C,D**) infected mice. Perisomatic nerve terminals are pseudocolored and shown in the insets in **A’-D’** and **A’’-D’’**. Scale bar in **A** = 7 µm for **A-D** and in **A’** = 850 nm for **A’-D’** and **A’’-D’’**. **E-H.** SBFSEM micrographs of neurons in layer V of the neocortex of mock-(**E,F**) and ME49-(**G,H**) infected mice. Perisomatic nerve terminals are pseudocolored and shown in the insets in **E’-H’** and **E’’-H’’**. Scale bar in **E** = 5 µm for **E-H** and in **E**’ = 150 nm for **E’-H’** and **E’’-H’’**. **I.** Quantification of perisomatic nerve terminals on entire neuronal somata in stratum pyramidalis of the hippocampus of mock- and ME49-infected mice. Each data point represents the total perisomatic synapses on one cell. Lines represent means +/− SEM. **** indicates p<0.0001 by Student’s t-test (n= 9). **J.** Quantification of cell thickness in stratum pyramidalis of mock- and ME49-infected hippocampi. Each data point represents the number of serial sections per cell. Lines represent means +/− SEM. ** indicates p<0.01 by Student’s t-test (n= 9). **K.** Despite reduced neuronal thickness in ME49-infected tissue, the ratio of perisomatic nerve terminal number to soma thickness is significantly reduced in parasite infected hippocampi. Each data point represents the ratio of perisomatic synapses to somata thickness per cell. Lines represent means +/− SEM. ** indicates p<0.01 by Student’s t-test (n= 9). **L**. The ratio of perisomatic nerve terminal number to soma thickness is significantly reduced in layer V of ME49-infected neocortex. Each data point represents the ratio of perisomatic synapses to somata thickness per cell. Lines represent means +/− SEM. ** indicates p<0.01 by Student’s t-test (n=7)

Similarly, it appeared as though fewer perisomatic synapses were present on neocortical neurons following *Toxoplasma* infection (Figure 3E-H). However, while the high density of neurons in stratum pyramidalis ensured we were able to identify neurons whose entire somas fell within the ~500 section SBFSEM datasets, the same was not true in neocortex. The sparser distribution of neurons in layer V of neocortex and the large size of the neurons meant few, if any, neurons could be analyzed in their entirety by SBFSEM. For this reason, we quantified the ratio of perisomatic synapses to the number of sections for neurons whose somas were mostly captured in the SBFSEM datasets (which we assessed based upon their entire nucleus being present in the dataset; for example, see Figure 4). As was the case in hippocampus, this revealed a significant reduction in the density of perisomatic synapses in neocortex following persistent *Toxoplasma* infection (Figure 3L). Together with GAD67-immunoreactivity data, these results reveal a significant loss of GABAergic perisomatic synapse following *Toxoplasma* infection.

**Figure 4.**
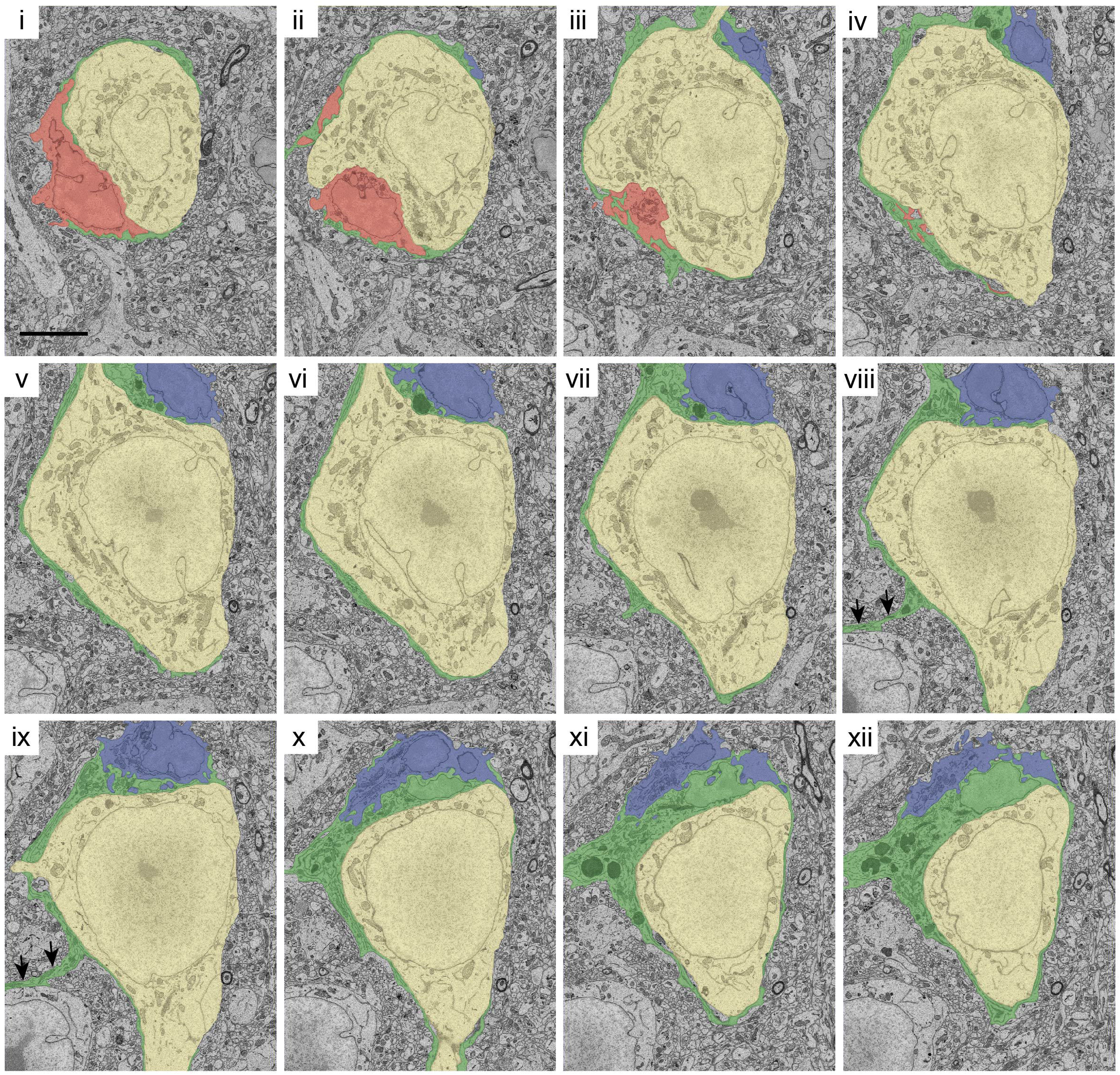
Phagocytic cells ensheath neuronal somata in *Toxoplasma*-infected neocortex. **A.** Twelve SBFSEM micrographs through the same pyramidal neuron (pseudocolored yellow) in layer V of ME49-infected neocortex. Three phagocytic cells ensheathing this neuron are pseudocolored blue, green, and red. Scale bar = 2.5 µm for all panels.

### Neuronal ensheathment by microglia following Toxoplasma infection

In assessing perisomatic synapses in SBFSEM datasets from *Toxoplasma*-infected tissue, we found that the somata of neurons were contacted and ensheathed by non-neuronal cells (something we failed to observe in mock-infected datasets). Ultrastructural analysis of these ensheathing cells revealed they displayed long and intricate processes, relatively dark cytoplasm, lysosomal dense granules, prominent Golgi apparatus, and phagosomes (Figure 2C, Supplemental Figure 2, Supplemental Movie) – all morphological features of phagocytes (i.e. resident microglia or derived from infiltrating monocytes) (Tremblay et al. 2010; Chen et al. 2014; Savage et al. 2018; Yamasaki et al. 2014). Ensheathment of neuronal somas by these phagocytes (in both neocortex and hippocampus of *Toxoplasma*-infected mice) was extensive and in several cases, multiple phagocytes ensheathed a single neuron. Figure 4 shows such an example from ME49-infected neocortex in which a single neuron is ensheathed by three distinct phagocytes. In many cases, little somal surface of neurons remains free of phagocyte-contact, and thus, little space of the neuronal cell surface remains available for contact by perisomatic nerve terminals. Remarkably, an individual phagocyte’s reach was not limited to a single neuronal soma but each often ensheathed portions of multiple somas (Figure 4).

While SBFSEM datasets provide a wealth of ultrastructural data, they are limited in scope and these observations could represent rare events that we were fortunate to capture in these datasets. To assess larger regions of hippocampus and neocortex for phagocyte-neuron interactions, we immunolabeled mock- and ME49-infected tissue with antibodies against Ionizing Calcium Binding Adapter 1 (Iba1) which is expressed by resident microglia in the healthy brain and by both reactive microglia and macrophages derived from infiltrating monocytes, following injury, infection, or inflammation (Yamasaki et al. 2014). In mock-infected mice, Iba1^+^ cells exhibited stellate morphologies, a hallmark of surveillant microglial phenotypes. Moreover, few Iba1^+^ resident microglia were observed in SP of CA1 in control mice (Figure 5). In contrast, in ME49-infected mice, Iba1^+^ phagocytes resembled the morphology of reactive microglia (i.e. increased cell size and thicker, shorter processes; Ransohoff and Perry 2019) and were abundant in SP of CA1 and all other regions of hippocampus and neocortex (Figure 5). Moreover, this analysis showed widespread Iba1^+^ cells in close apposition with other brain cells (labeled either by their nuclei [DAPI] or with NeuroTrace) in *Toxoplasma*-infected neocortex and hippocampus (Figure 5A-D, G-J). We quantified the number of Iba1^+^ cells that were surrounding or contacting Iba1^−^/DAPI^+^ cells (see arrows in Figure 5B,H). This quantification confirmed widespread, significant contact and ensheathment of neurons by Iba1^+^ cells in the hippocampus and neocortex of ME49-infected brains (Figure 5M,N).

**Figure 5.**
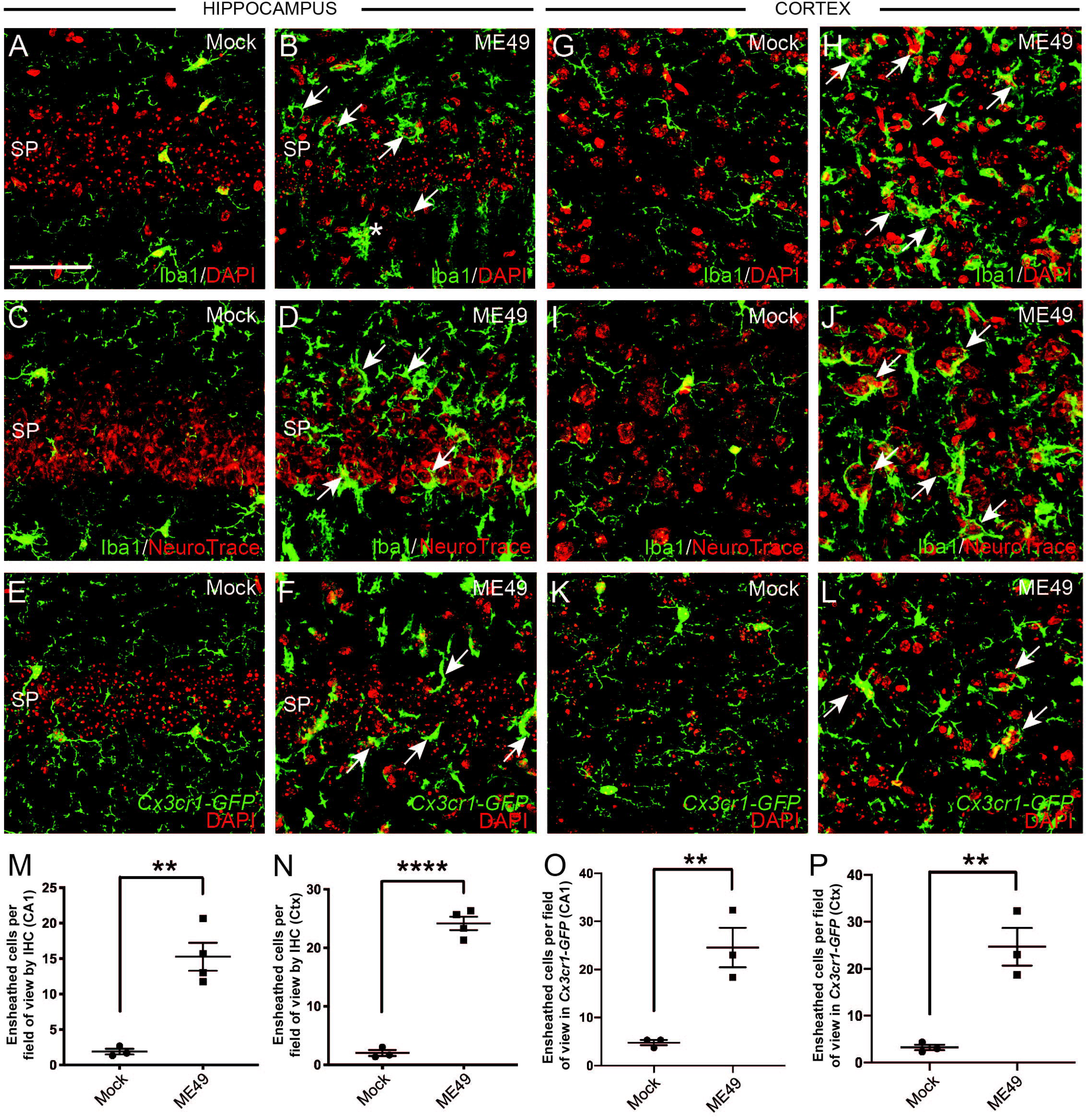
Widespread ensheathment of cells by microglia in the hippocampus and neocortex of *Toxoplasma*-infected mice. **A,B.** Immunolabeling of microglia with antibodies against Iba1 and labeling of nuclei with DAPI in mock-(**A**) and ME49-(**B**) infected CA1 of hippocampus. SP denotes stratum pyramidalis. Arrows highlight cells ensheathed by Iba1^+^ cells. Scale bar = 50 µm for A-L. **C,D.** Iba1-immunolabeling and cell labeling with NeuroTrace in mock-(**C**) and ME49-(**D**) infected CA1 of hippocampus. Arrows highlight cells ensheathed by Iba1^+^ cells in ME49-infected tissue. **E,F.** Genetic labeling of phagocytic cells in mock-(**E**) and ME49-(**F**) infected CA1 of *Cx3cr1-GFP* mice. Arrows highlight cells ensheathed by GFP^+^ cells in ME49-infected tissue. **G,H.** Immunolabeling of microglia with antibodies against Iba1 and labeling of nuclei with DAPI in mock-(**G**) and ME49-(**H**) infected layer V of neocortex. Arrows highlight cells ensheathed by Iba1^+^ cells. **I,J.** Iba1-immunolabeling and cell labeling with NeuroTrace in mock-(**I**) and ME49-(**J**) infected layer V of neocortex. Arrows highlight cells ensheathed by Iba1^+^ cells in ME49-infected tissue. **K,L.** Genetic labeling of phagocytic cells in mock-(**K**) and ME49-(**L**) infected neocortex of *Cx3cr1-GFP* mice. Arrows highlight cells ensheathed by GFP^+^ cells in ME49-infected tissue. **M,N.** Quantification of the number of DAPI^+^ cell somas ensheathed by Iba1^+^ cells in stratum pyramidalis of CA1 (**M**) or layer V of neocortex (**N**). Lines represent means +/− SEM. ** indicates p<0.01 and **** indicates p<0.0001 by Student’s t-test (n=4). **O,P**. Quantification of the number of DAPI^+^ cell somas ensheathed by GFP^+^ cells in stratum pyramidalis of CA1 (**O**) or layer V of neocortex (**P**). Lines represent means +/− SEM. ** indicates p<0.01 by Student’s t-test (n=3).

Since Iba1-immunoreactivity alone cannot distinguish reactive resident microglia from macrophages derived from infiltrating monocytes, we assessed phagocytic ensheathment of neurons in *Cx3cr1-GFP* mice infected with *Toxoplasma gondii*. Microglia, but not infiltrating monocytes, have been reported to express high levels of GFP in inflamed brains of these transgenic reporter mice (Yamasaki et al. 2014). In mock-infected *Cx3cr1-GFP*, GFP^+^ cells exhibited characteristic surveillant microglia morphology in all brain regions and few GFP^+^ cells were present in SR of CA1 (Figure 5,6). In contrast, GFP^+^ microglia appeared strikingly different in infected mice. First, in SP of CA1 and layer V of neocortex, microglia appeared reactive, with larger cell bodies and thicker, shorter processes following infection (Figure 5,6 and Supplemental Figure 2). Second, GFP^+^ microglia appeared strikingly different in distinct hippocampal layers of infected *Cx3cr1-GFP* mice, with rod-like microglia being present in stratum radiatum (SR) and stratum lacunose-moleculare (SLM) (Supplemental Figure 2). Similar rod-like, Iba1^+^ cells were observed in SR of *Toxoplasma*-infected mice (see asterisk in Figure 5B) (also see Wyatt-Johnson et al. 2017). Finally, and most importantly, GFP^+^ microglia ensheathed GFP^−^ cells in ME49-infected hippocampus and cortex (Figure 5O, P). These data, coupled with the ultrastructural morphology of these cells in SBFSEM datasets (Figure 4, Supplemental Figure 2, and Supplemental Movie) (Yamasaki et al. 2014), suggest that a significant fraction of the neuron-ensheathing phagocytes in *Toxoplasma-*infected brains are reactive microglia.

**Figure 6.**
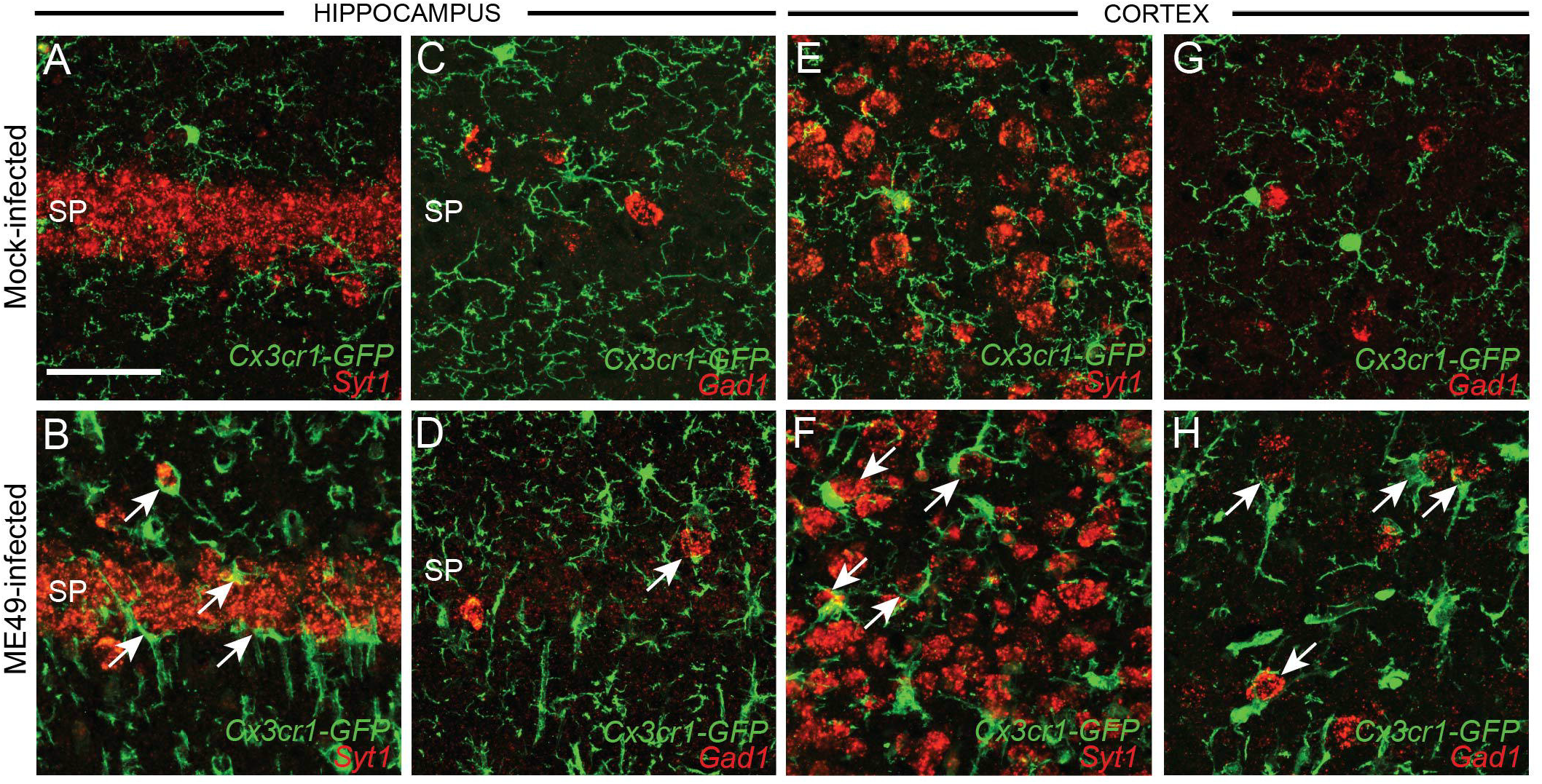
Microglia ensheath both excitatory and inhibitory neurons in parasite-infected hippocampus and neocortex. **A-D**. *In situ* hybridization for *Syt1* mRNA and *Gad1* mRNA to label excitatory and inhibitory neurons, respectively, in CA1 of mock- or ME49-infected *Cx3cr1-GFP* mice. Arrows highlight *Syt1*^+^ and *Gad1*^*+*^ cells ensheathed by GFP^+^ phagocytic cells. **E-H**. *In situ* hybridization for *Syt1* mRNA and *Gad1* mRNA to label excitatory and inhibitory neurons, respectively, in cortical layer V of mock- or ME49-infected *Cx3cr1-GFP* mice. Arrows highlight *Syt1*^+^ and *Gad1*^*+*^ cells ensheathed by GFP^+^ phagocytic cells. Scale bar in **A** = 50 µm for all panels.

Finally, we sought to determine whether reactive microglia ensheath both excitatory and inhibitory neurons following persistent *Toxoplasma* infection. As described above, we used *in situ* hybridization for *Syt1* and *Gad1* mRNA to label these populations of neurons. *In situ* hybridization in *Cx3cr1-GFP* mock- and ME49-infected tissue revealed that reactive GFP^+^ cells ensheath both excitatory and inhibitory neurons in hippocampus and neocortex (Figure 6).

### Widespread microglial removal of perisomatic synapses post chronic parasitic infection

Microglia have been implicated in synaptic remodeling both in development and various models of injury and disease (Kettenmann et al. 2013). Since immunolabeling and ultrastructural data revealed inhibitory perisomatic synapse loss and microglial activation following persistent *Toxoplasma gondii* infection, it seemed plausible that these reactive microglia may be involved in stripping inhibitory perisomatic nerve terminals. To begin to test this, several approaches were employed. First, we used SBFSEM datasets to demonstrate that microglia contact, surround, and potentially displace perisomatic synapses (Figure 7A-C). Similar observations in SBFSEM datasets in other models of inflammation have been suggested to support synaptic stripping by microglia (Chen et al. 2014). Moreover, SBFSEM images illustrated frequent examples of phagosomes in microglial processes adjacent to neuronal somata in ME49-infected mice (Figure 7B, 8A,B). Finally, we employed immunohistochemistry against Cluster of Differentiation 68 (CD68), a phagocytic cell marker, to assess the phagocytic nature of Iba1^+^ and GFP^+^ cells following parasitic infection. Enhanced CD68-immunoreactivity was observed in hippocampus and cortex of *Toxoplasma gondii*-infected brains, compared with mock-infected controls (Figure 8C-J). Furthermore, colocalization of CD68 with Iba1^+^ phagocytes and GFP^+^ microglia (in *Cx3cr1-GFP* mice) confirmed that microglia in ME49-infected tissue were not only reactive, but exhibited enhanced phagocytic activity compared to those in controls (Figure 8C-J). Taken together, these three sets of results strongly suggest reactive microglia participate in synaptic stripping following persistent *Toxoplasma gondii* infection.

**Figure 7.**
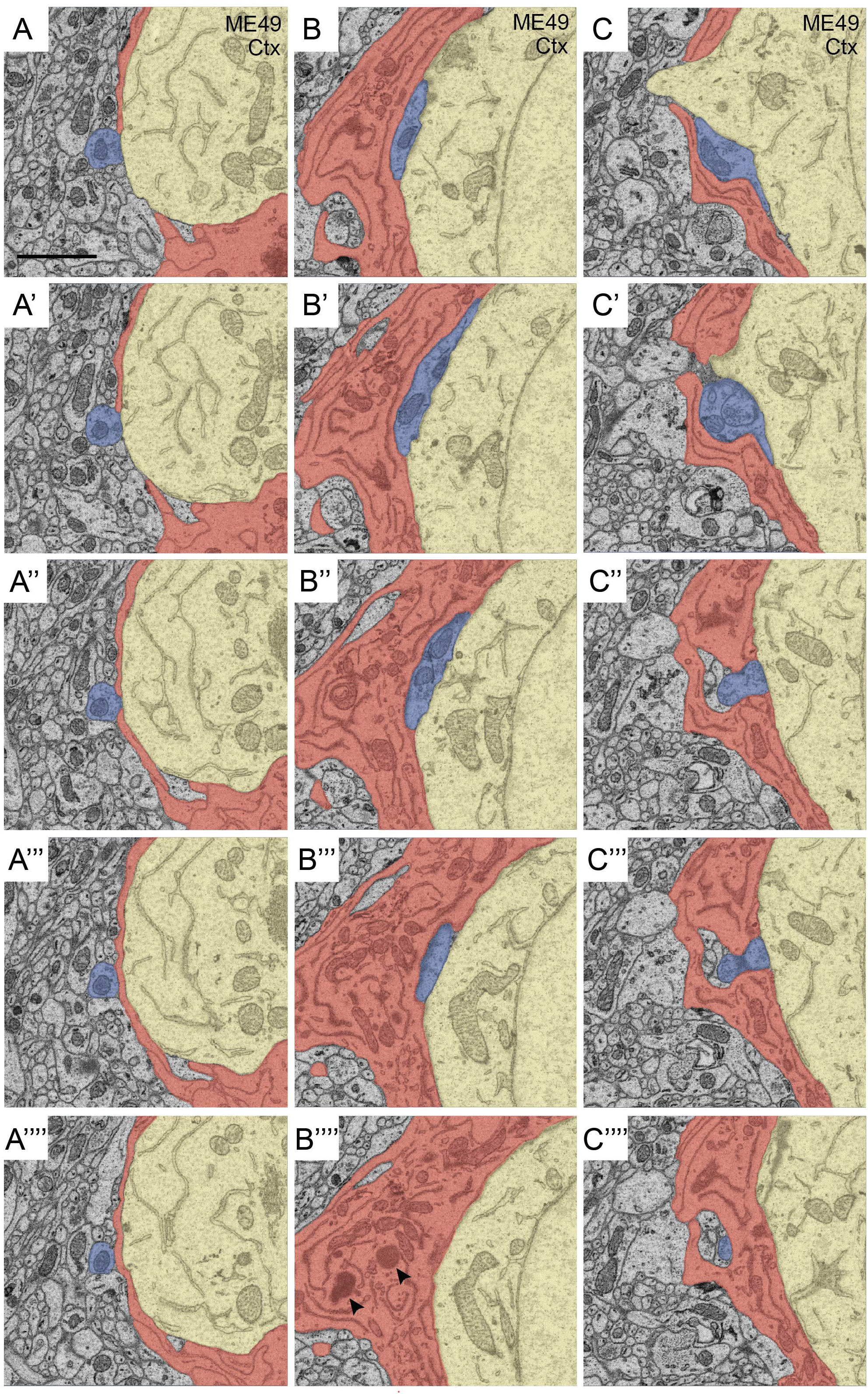
Perisomatic synapses are ensheathed by microglia in *Toxoplasma*-infected brain tissue. **A-C.** Examples of 3 perisomatic nerve terminals (in blue) ensheathed by phagocytic microglia (in red) on a neocortical neuron (yellow). 5 micrographs shown for each example from this SBFSEM dataset. Scale bar in **A** = 1 µm for all panels.

**Figure 8.**
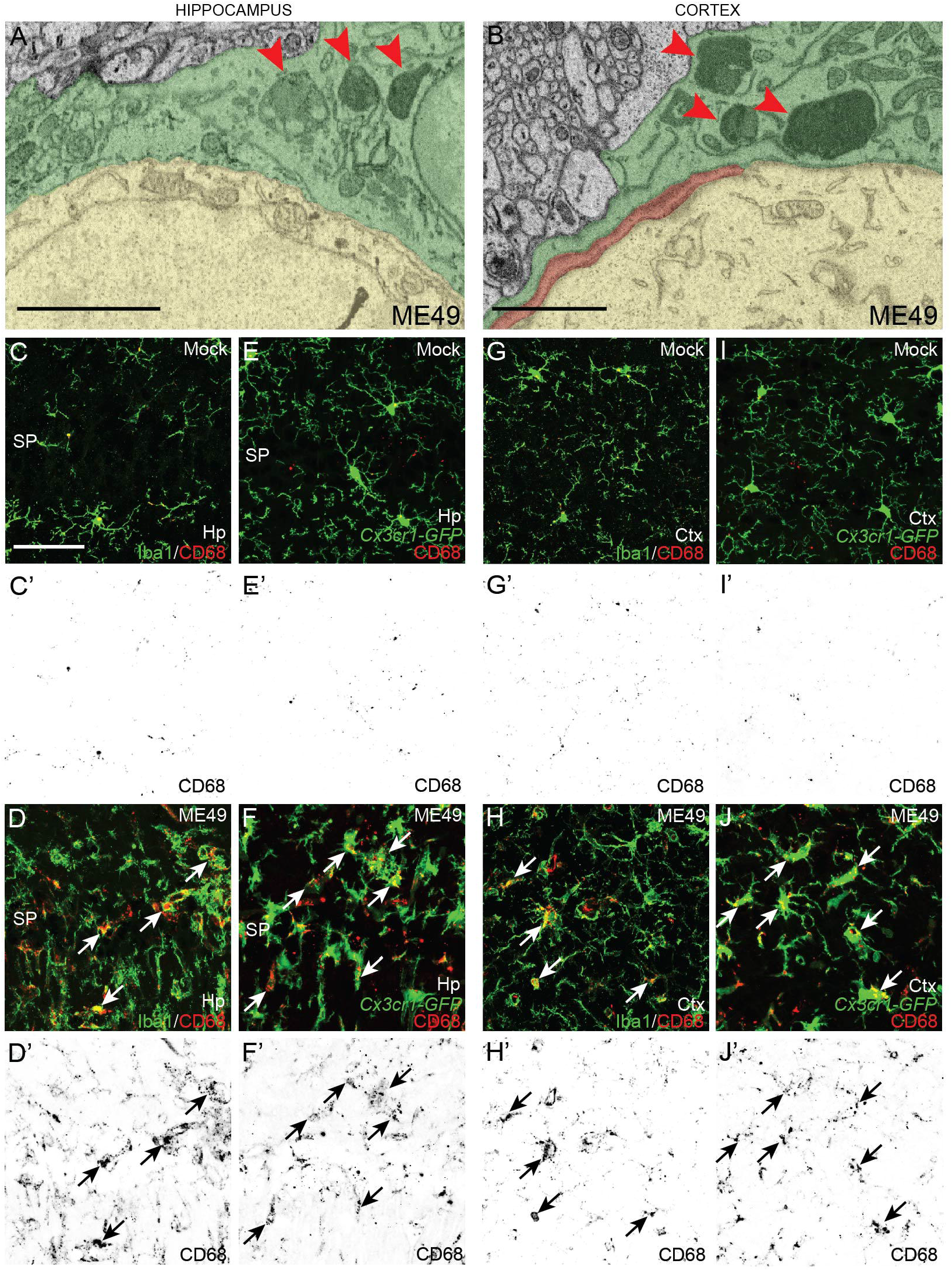
Phagocytic microglia ensheath neurons in *Toxoplasma*-infected brain tissue. **A,B.** SBFSEM micrographs show phagosomes within the microglial ensheathing neuronal cell somas in ME49-infected hippocampus (**A**) and cortex (**B**). Phagosomes highlighted by red arrowheads. Neuronal somas pseudocolored in yellow. Microglia pseudocolored in green and red. **C-J**. Immunolabeling for CD68 in stratum pyramidalis (SP) of CA1 and layer V of neocortex in mock- or ME49-infected *Cx3cr1-GFP* mice. Arrows highlight microglia with elevated levels of CD68, a marker of phagocytosis, in ME49-infected cortex and hippocampus. Scale bar in **A** = 1 µm; in **B** = 1.25 µm; in **C** = 50 µm for **C-J**.

## DISCUSSION

### Synaptic changes following parasitic infection

Here, we report the novel discovery that inhibitory perisomatic synapses are lost following persistent infection with *Toxoplasma gondii*. In. hippocampus and neocortex these perisomatic synapses are generated by Parv-expressing, fast-spiking interneurons (Kawaguchi 1993; Kawaguchi and Kubota 1998,1997; Pi et al. 2013; Pfeffer et al. 2013). Although Parv^+^ interneurons represent only a fraction of the total neurons in hippocampus or neocortex (<10%)(Freund and Buzsaki 1996; Tremblay et al. 2016), the perisomatic synapses they generate on excitatory pyramidal neurons and other Parv^+^ interneurons (Pi et al. 2013) play essential roles in controlling the spread of neural activity (Hensch 2005), in regulating oscillatory network activity (Buzsaki and Wang 2012; Tukker et al. 2007; Bartos et al. 2002), and in regulating neural plasticity during development, learning and plasticity (Fagiolini et al. 2004; Donato et al. 2013; Kuhlman et al 2013). Disruption of Parv^+^ synapses (or Parv^+^ interneurons) in neocortex or hippocampus has been associated with epileptiform activity (Schwaller et al. 2004; Paz and Huguenard 2015; Su et al. 2016) and optogenetic stimulation of these interneurons at the onset of seizure activity is sufficient to halt electrographic seizures and reduce behavioral seizures (Krook-Magnuson et al. 2013). Thus, the loss of GABAergic perisomatic synapses reported here following persistent *Toxoplasma* infection is likely to underlie spontaneous seizures and enhanced susceptibility to drug-induced seizures observed in infected mice (Brooks et al. 2015; David et al. 2016). It is important to point out, however, that alternative mechanisms underlying epileptiform activity following chronic *Toxoplasma gondii* infection have been proposed, including the downregulation of astrocyte-derived excitatory amino acid transporter 2 (EAAT2, also called Glt1), which is essential for clearing extracellular glutamate from the synaptic cleft following neurotransmission (David et al. 2016; Wohlfert et al. 2017). Together, the loss of both perisomatic inhibitory synapses and elevated levels of extracellular glutamate (due to the loss of astrocyte-derived EAAT2; David et al. 2016) offer a two-hit model for how seizures develop in patients suffering from Toxoplasmic encephalitis.

In addition to seizures, a large number of studies, including a recent case-control study with over 80,000 individuals, have revealed a strong association between *Toxoplasma gondii* infection and neuropsychiatric disorders, including schizophrenia (Dickenson et al. 2007, 2014; Flegr and Horacek 2018; Burgdorf et al. 2019). Given that *Toxoplasma* parasites are known to increase dopamine metabolism in their hosts and even generate dopamine themselves (Prandovszky et al. 2011; Martin et al. 2015; Ihara et al. 2016; Gaskell et al. 2009) and neuropsychiatric illnesses have long been attributed to dopamine dysfunction (Brisch et al. 2014), it has been hypothesized that the association between persistent *Toxoplasma* infection and schizophrenia is due to altered dopamine signaling (Skallova et al. 2006; Elsheikha et al. 2016; Stock et al. 2017; but see also Wang et al. 2015, 2017). However, it is noteworthy that the impairment of Parv^+^ interneurons and GABAergic perisomatic synapses has also been strongly associated with the development of neuropsychiatric disorders, such as schizophrenia (Belforte et al. 2010; Marin 2012; Gonzalez-Burgos et al. 2010, 2011; Sgado et al. 2011; Gonzalez-Burgos and Lewis 2012; Lewis et al. 2012). Similarly, defects in Parv^+^ interneurons and synapses have been reported to promote the onset of epilepsy (Jiang et al. 2016), which has been reported to be a risk factor in *Toxoplasma*-infected individuals (Ngoungou et al. 2015; Palmer 2007). Thus, results presented here offer an alternative mechanism by which persistent *Toxoplasma* infection may contribute to the development of neuropsychiatric and neurological illnesses.

Finally, while we focus on the loss of perisomatic synapses in this study, it is likely that far more inhibitory synapses are affected or lost following parasitic infection than just perisomatic synapses formed by Parv^+^ interneurons. Thus, *Toxoplasma gondii* infection likely has a far wider influence on inhibitory synapses and neurotransmission than reported here. This hypothesis stems from the observation that persistent ME49 infection results in altered GAD67 immunoreactivity (i.e. the loss of its punctate appearance) in regions that lack these Parv^+^ perisomatic inhibitory synapses. This raises the question of what widespread redistribution of GAD67 following persistent *Toxoplasma* infection might reflect. Data presented here suggests that the loss of punctate GAD67-immunoreactivity reflect the loss of inhibitory synapses. But what about the increased and diffuse nature of GAD67-immunoreactivity outside of these synaptic zones (Brooks et al. 2015)? One possibility is that GAD67 may be upregulated by other cells following parasitic infection. In fact, one recent study suggested that *Toxoplasma* infection induces an upregulation of GAD67 by microglia (Bhandage et al. 2019). Unfortunately, our *in situ* hybridization studies do not support expression of *Gad1* mRNA by hippocampal or neocortical microglia following persistent ME49 infection (Figure 6B,D,F,H). Instead, we suspect that the diffuse localization of GAD67 immunoreactivity following ME49 infection indicates reduced levels of GAD67 from remaining inhibitory nerve terminals and its increase in other compartments of the neurons (i.e. neurites and somata). During development, components of the presynaptic machinery, including GAD67, appear diffusely localized throughout the neuron, and only become concentrated at the presynaptic terminal following the process of synaptic differentiation and maturation (Fox and Umemori 2006; Fox et al. 2007). Since inhibitory interneurons exhibit plasticity following activity, disease or injury – including the ability to integrate into new circuits in the adult brain (Baraban et al. 2009; Donato et al. 2013; Tang et al. 2014; Mukherjee et al. 2019) – the redistribution of GAD67 following persistent *Toxoplasma* infection may reflect the re-initiation of a developmental program to promote inhibitory synapse formation or plasticity. As such, the infected brain may be undergoing cycles of stripping and regenerating inhibitory synapses as a result of inflammation or the parasite itself.

### Loss of perisomatic synapses coincides with microglial ensheathment of neuronal somata in parasite-infected brains

While our initial intent was to assess perisomatic synapse density following parasite-infection using a state-of-the-art ultrastructural imaging modality (SBFSEM), we unexpectedly discovered that non-neuronal, phagocytic cells ensheathed the cell bodies of neurons in both hippocampus and neocortex of infected mice. The electron dense cytoplasm of these process-bearing cells, the presence of lysosomes and phagosomes, the relatively sparse volume of cytoplasm, and the distribution of electron dense heterochromatin along the nuclear membrane all suggest these are microglia (Savage et al. 2018). This suggestion was supported by results demonstrating Iba1^+^ and *Cx3cr1-GFP*^+^ cells are closely apposed to excitatory and inhibitory neurons following persistent parasitic infection. The association of reactive, phagocytic microglia with neuronal somata is not unique to animals with latent *Toxoplasma gondii* infection, as these cellular interactions have been observed in a number of neurodegenerative and neuroimmune disorders (Haga et al. 1989; Peterson et al. 2001; Neumann et al. 2006). While results presented here implicate microglia in neuronal ensheathment (and potentially synaptic loss), they do not rule out roles for macrophages derived from infiltrating monocytes which are notoriously difficult to unambiguously differentiate from resident microglia following injury, infection, or inflammation (Koeniger and Kuerlen 2017; but see Yamasaki et al. 2014). In fact, although CD68-immunoreactivity is often used as a marker of microglial cells, our results identified numerous GFP^−^/CD68^+^ cells in ME49-infected *Cx3cr1-GFP* mice which may represent phagocytic infiltrating monocytes in neocortex and hippocampus of parasite-infected brains.

Why might microglia ensheath neuronal cell bodies following parasitic infection? Over half a century ago, microglia were observed ensheathing lower motoneuron somas following axotomy and this coincided with the loss of perisomatic synapses onto these cells (Blinzinger and Kreutzberg 1968). It was hypothesized that these microglia played an active role in the removal of these synapses, a process termed synaptic stripping. Synaptic stripping has been observed not only after axotomy or injury, but also following infection or inflammation (Di Liberto et al. 2018; Kreutzfeldt et al. 2013; Trapp et al. 2007; Chen et al. 2014), and it involves not only excitatory perisomatic synapses (like those on lower motoneurons; Blinzinger and Kreutzberg 1968) but also inhibitory perisomatic synapses (like those in neocortex and hippocampus; Di Liberto et al. 2018; Chen et al. 2014). While activation of microglia following insult and inflammation may lead to synaptic stripping, it does not in all cases (Siskova et al. 2009; Perry and O’Conner 2010), nor is activation a requirement for microglial to phagocytose synaptic elements. During development, non-reactive microglia remove supernumerary excitatory synapses (or reshape excitatory nerve terminals through trogocytosis) in the thalamus, neocortex and hippocampus (Tremblay et al. 2010; Schafer et al. 2012; Weinhard et al. 2018).

*Toxoplasma* infection is known to induce microglial activation (Dellacasa-Lindberg et al. 2011; Li et al. 2019), although it is unclear whether this is the result of direct parasite-derived signals, direct neuron-derived signals (Di Liberto et al. 2018), altered oscillatory network activity (Iaccarino et al. 2016), or other indirect pathways (such as inflammation). Here, we show that these reactive microglia not only ensheath excitatory and inhibitory neurons, but enwrap, and potentially phagocytose, perisomatic inhibitory synapses. Although we previously failed to observe changes in excitatory nerve terminals following parasitic infection (Brooks et al. 2015), other groups have reported altered connectivity, reduced levels of excitatory synaptic machinery, and reduced dendritic spine density (David et al. 2016; Mitra et al. 2013; Lang et al. 2018; Parlog et al. 2014; Wang et al. 2018), all suggesting these reactive microglial may phagocytose more than just inhibitory synapses following persistent parasite infection. The unique morphologies we observed for reactive microglia in different sublamina of the hippocampus, where excitatory and inhibitory synapses are largely restricted to non-overlapping regions, certainly support roles for these microglia beyond phagocytosing perisomatic inhibitory synapses following *Toxoplasma gondii* infection.

The ability of microglia to strip or prune synaptic elements has been proposed to be both pathological (as in Alzheimer’s Disease, Amyotrophic Lateral Sclerosis, and Lupus; Hong et al. 2016; Shi et al. 2017; Paolicelli et al. 2017; Krasemann et al. 2017; Bialas et al. 2017) and neuroprotective (Chen et al. 2014). At present it remains unclear whether the loss of perisomatic synapses following *Toxoplasma gondii* infection is pathological (and the direct cause of epileptiform activity) or beneficial (and may be a homeostatic response to other alterations following infection or inflammation). Insight into this question may come from assessing molecular signatures of the populations of microglia present in the parasite-infected brain. In other conditions, such analysis has shed light on whether microglia are pro- or anti-inflammatory, scavenging debris, or promoting tissue homeostasis (Butovsky and Weiner 2018). In cases where microglia have been implicated in the pathological loss of synapses, it appears to be mediated through the complement pathway (Hong et al. 2016; Sekar et al. 2016; Vasek et al. 2016; Hammond et al. BioRxiv), a similar pathway used to prune the excess of excitatory synapses during development (Schafer et al. 2012; Stevens et al. 2007). In contrast, the proposed neuroprotective role for microglial synaptic stripping appear to be complement-independent (Chen et al. 2014; Di Liberto et al. 2018). While future studies will be needed to resolve the beneficial or pathological consequences of synapse loss reported here, recent studies showing activation of the complement pathway in microglia following *Toxoplasma gondii* infection (Li et al. 2019; Xiao et al. 2016) suggest that the actions of microglia in this case may be pathological.

## ACKNOWLEDGEMENTS

This work was supported in part by the National Institutes of Health – AI124677 (M.A.F., I.J.B.) and NS105141 (M.A.F.). We thank Dr. M.Theus (Virginia Tech) for generously supplying *Cx3Cr1-GFP* mice and for Mr. U.Sabbagh for comments on the manuscript.

## REFERENCES

Alsaady, I., Tedford, E., Alsaad, M., Bristow, G., Kohli, S., Murray, M., Reeves, M., Vijayabaskar, M.S., Clapcote, S.J., Wastling, J., et al. (2019). Downregulation of the Central Noradrenergic System by Toxoplasma gondii Infection. Infect Immun 87.

Baraban, S.C., Southwell, D.G., Estrada, R.C., Jones, D.L., Sebe, J.Y., Alfaro-Cervello, C., Garcia-Verdugo, J.M., Rubenstein, J.L., and Alvarez-Buylla, A. (2009). Reduction of seizures by transplantation of cortical GABAergic interneuron precursors into Kv1.1 mutant mice. Proc Natl Acad Sci U S A 106, 15472–15477.

Bartos, M., Vida, I., Frotscher, M., Meyer, A., Monyer, H., Geiger, J.R., and Jonas, P. (2002). Fast synaptic inhibition promotes synchronized gamma oscillations in hippocampal interneuron networks. Proc Natl Acad Sci U S A 99, 13222–13227.

Belforte, J.E., Zsiros, V., Sklar, E.R., Jiang, Z., Yu, G., Li, Y., Quinlan, E.M., and Nakazawa, K. (2010). Postnatal NMDA receptor ablation in corticolimbic interneurons confers schizophrenia-like phenotypes. Nat Neurosci 13, 76–83.

Berdoy, M., Webster, J.P., and Macdonald, D.W. (2000). Fatal attraction in rats infected with Toxoplasma gondii. Proc Biol Sci 267, 1591–1594.

Beste, C., Getzmann, S., Gajewski, P.D., Golka, K., and Falkenstein, M. (2014). Latent Toxoplasma gondii infection leads to deficits in goal-directed behavior in healthy elderly. Neurobiol Aging 35, 1037–1044.

Bhandage, A.K., Kanatani, S., and Barragan, A. (2019). Toxoplasma-Induced Hypermigration of Primary Cortical Microglia Implicates GABAergic Signaling. Front Cell Infect Microbiol 9, 73.

Bialas, A.R., Presumey, J., Das, A., van der Poel, C.E., Lapchak, P.H., Mesin, L., Victora, G., Tsokos, G.C., Mawrin, C., Herbst, R., et al. (2017). Microglia-dependent synapse loss in type I interferon-mediated lupus. Nature 546, 539–543.

Birnbaum, R., and Weinberger, D.R. (2017). Genetic insights into the neurodevelopmental origins of schizophrenia. Nat Rev Neurosci 18, 727–740.

Blinzinger, K., and Kreutzberg, G. (1968). Displacement of synaptic terminals from regenerating motoneurons by microglial cells. Z Zellforsch Mikrosk Anat 85, 145–157.

Brisch, R., Saniotis, A., Wolf, R., Bielau, H., Bernstein, H.G., Steiner, J., Bogerts, B., Braun, K., Jankowski, Z., Kumaratilake, J., et al. (2014). The role of dopamine in schizophrenia from a neurobiological and evolutionary perspective: old fashioned, but still in vogue. Front Psychiatry 5, 47.

Brooks, J.M., Carrillo, G.L., Su, J., Lindsay, D.S., Fox, M.A., and Blader, I.J. (2015). Toxoplasma gondii Infections Alter GABAergic Synapses and Signaling in the Central Nervous System. MBio 6, e01428–01415.

Burgdorf, K.S., Trabjerg, B.B., Pedersen, M.G., Nissen, J., Banasik, K., Pedersen, O.B., Sorensen, E., Nielsen, K.R., Larsen, M.H., Erikstrup, C., et al. (2019). Large-scale study of Toxoplasma and Cytomegalovirus shows an association between infection and serious psychiatric disorders. Brain Behav Immun 79, 152–158.

Butovsky, O., and Weiner, H.L. (2018). Microglial signatures and their role in health and disease. Nat Rev Neurosci 19, 622–635.

Buzsaki, G., and Wang, X.J. (2012). Mechanisms of gamma oscillations. Annu Rev Neurosci 35, 203–225.

Cabral, C.M., Tuladhar, S., Dietrich, H.K., Nguyen, E., MacDonald, W.R., Trivedi, T., Devineni, A., and Koshy, A.A. (2016). Neurons are the Primary Target Cell for the Brain-Tropic Intracellular Parasite Toxoplasma gondii. PLoS Pathog 12, e1005447.

Cardona, A., Saalfeld, S., Schindelin, J., Arganda-Carreras, I., Preibisch, S., Longair, M., Tomancak, P., Hartenstein, V., and Douglas, R.J. (2012). TrakEM2 software for neural circuit reconstruction. PLoS One 7, e38011.

Chen, Z., Jalabi, W., Hu, W., Park, H.J., Gale, J.T., Kidd, G.J., Bernatowicz, R., Gossman, Z.C., Chen, J.T., Dutta, R., et al. (2014). Microglial displacement of inhibitory synapses provides neuroprotection in the adult brain. Nat Commun 5, 4486.

Coombes, J.L., Charsar, B.A., Han, S.J., Halkias, J., Chan, S.W., Koshy, A.A., Striepen, B., and Robey, E.A. (2013). Motile invaded neutrophils in the small intestine of Toxoplasma gondii-infected mice reveal a potential mechanism for parasite spread. Proc Natl Acad Sci U S A 110, E1913–1922.

Courret, N., Darche, S., Sonigo, P., Milon, G., Buzoni-Gatel, D., and Tardieux, I. (2006). CD11c- and CD11b-expressing mouse leukocytes transport single Toxoplasma gondii tachyzoites to the brain. Blood 107, 309–316.

David, C.N., Frias, E.S., Szu, J.I., Vieira, P.A., Hubbard, J.A., Lovelace, J., Michael, M., Worth, D., McGovern, K.E., Ethell, I.M., et al. (2016). GLT-1-Dependent Disruption of CNS Glutamate Homeostasis and Neuronal Function by the Protozoan Parasite Toxoplasma gondii. PLoS Pathog 12, e1005643.

Dellacasa-Lindberg, I., Fuks, J.M., Arrighi, R.B., Lambert, H., Wallin, R.P., Chambers, B.J., and Barragan, A. (2011). Migratory activation of primary cortical microglia upon infection with Toxoplasma gondii. Infect Immun 79, 3046–3052.

Denk, W., and Horstmann, H. (2004). Serial block-face scanning electron microscopy to reconstruct three-dimensional tissue nanostructure. PLoS Biol 2, e329.

Di Liberto, G., Pantelyushin, S., Kreutzfeldt, M., Page, N., Musardo, S., Coras, R., Steinbach, K., Vincenti, I., Klimek, B., Lingner, T., et al. (2018). Neurons under T Cell Attack Coordinate Phagocyte-Mediated Synaptic Stripping. Cell 175, 458–471 e419.

Dickerson, F., Origoni, A., Schweinfurth, L.A.B., Stallings, C., Savage, C.L.G., Sweeney, K., Katsafanas, E., Wilcox, H.C., Khushalani, S., and Yolken, R. (2018). Clinical and Serological Predictors of Suicide in Schizophrenia and Major Mood Disorders. J Nerv Ment Dis 206, 173–178.

Dickerson, F., Stallings, C., Origoni, A., Schroeder, J., Khushalani, S., and Yolken, R. (2014). Mortality in schizophrenia: clinical and serological predictors. Schizophr Bull 40, 796–803.

Dickerson, F., Wilcox, H.C., Adamos, M., Katsafanas, E., Khushalani, S., Origoni, A., Savage, C., Schweinfurth, L., Stallings, C., Sweeney, K., et al. (2017). Suicide attempts and markers of immune response in individuals with serious mental illness. J Psychiatr Res 87, 37–43.

Donato, F., Rompani, S.B., and Caroni, P. (2013). Parvalbumin-expressing basket-cell network plasticity induced by experience regulates adult learning. Nature 504, 272–276.

Elsheikha, H.M., Busselberg, D., and Zhu, X.Q. (2016). The known and missing links between Toxoplasma gondii and schizophrenia. Metab Brain Dis 31, 749–759.

Fagiolini, M., Fritschy, J.M., Low, K., Mohler, H., Rudolph, U., and Hensch, T.K. (2004). Specific GABAA circuits for visual cortical plasticity. Science 303, 1681–1683.

Flegr, J., and Horacek, J. (2018). Toxoplasmosis, but not borreliosis, is associated with psychiatric disorders and symptoms. Schizophr Res 197, 603–604.

Fox, M.A., and Umemori, H. (2006). Seeking long-term relationship: axon and target communicate to organize synaptic differentiation. J Neurochem 97, 1215–1231.

Fox, M.A., Sanes, J.R., Borza, D.B., Eswarakumar, V.P., Fassler, R., Hudson, B.G., John, S.W., Ninomiya, Y., Pedchenko, V., Pfaff, S.L., et al. (2007). Distinct target-derived signals organize formation, maturation, and maintenance of motor nerve terminals. Cell 129, 179–193.

Freund, T.F., and Buzsaki, G. (1996). Interneurons of the hippocampus. Hippocampus 6, 347–470.

Gaskell, E.A., Smith, J.E., Pinney, J.W., Westhead, D.R., and McConkey, G.A. (2009). A unique dual activity amino acid hydroxylase in Toxoplasma gondii. PLoS One 4, e4801.

Gatkowska, J., Wieczorek, M., Dziadek, B., Dzitko, K., and Dlugonska, H. (2013). Sex-dependent neurotransmitter level changes in brains of Toxoplasma gondii infected mice. Exp Parasitol 133, 1–7.

Gonzalez-Burgos, G., Fish, K.N., and Lewis, D.A. (2011). GABA neuron alterations, cortical circuit dysfunction and cognitive deficits in schizophrenia. Neural Plast 2011, 723184.

Gonzalez-Burgos, G., Hashimoto, T., and Lewis, D.A. (2010). Alterations of cortical GABA neurons and network oscillations in schizophrenia. Curr Psychiatry Rep 12, 335–344.

Gonzalez-Burgos, G., and Lewis, D.A. (2012). NMDA receptor hypofunction, parvalbumin-positive neurons, and cortical gamma oscillations in schizophrenia. Schizophr Bull 38, 950–957.

Haga, S., Akai, K., and Ishii, T. (1989). Demonstration of microglial cells in and around senile (neuritic) plaques in the Alzheimer brain. An immunohistochemical study using a novel monoclonal antibody. Acta Neuropathol 77, 569–575.

Hamm, J.P., Peterka, D.S., Gogos, J.A., and Yuste, R. (2017). Altered Cortical Ensembles in Mouse Models of Schizophrenia. Neuron 94, 153–167 e158.

Hammer, S., Carrillo, G.L., Govindaiah, G., Monavarfeshani, A., Bircher, J.S., Su, J., Guido, W., and Fox, M.A. (2014). Nuclei-specific differences in nerve terminal distribution, morphology, and development in mouse visual thalamus. Neural Dev 9, 16.

Haroon, F., Handel, U., Angenstein, F., Goldschmidt, J., Kreutzmann, P., Lison, H., Fischer, K.D., Scheich, H., Wetzel, W., Schluter, D., et al. (2012). Toxoplasma gondii actively inhibits neuronal function in chronically infected mice. PLoS One 7, e35516.

Hensch, T.K. (2005). Critical period plasticity in local cortical circuits. Nat Rev Neurosci 6, 877–888.

Hong, S., Beja-Glasser, V.F., Nfonoyim, B.M., Frouin, A., Li, S., Ramakrishnan, S., Merry, K.M., Shi, Q., Rosenthal, A., Barres, B.A., et al. (2016). Complement and microglia mediate early synapse loss in Alzheimer mouse models. Science 352, 712–716.

Iaccarino, H.F., Singer, A.C., Martorell, A.J., Rudenko, A., Gao, F., Gillingham, T.Z., Mathys, H., Seo, J., Kritskiy, O., Abdurrob, F., et al. (2016). Gamma frequency entrainment attenuates amyloid load and modifies microglia. Nature 540, 230–235.

Ihara, F., Nishimura, M., Muroi, Y., Mahmoud, M.E., Yokoyama, N., Nagamune, K., and Nishikawa, Y. (2016). Toxoplasma gondii Infection in Mice Impairs Long-Term Fear Memory Consolidation through Dysfunction of the Cortex and Amygdala. Infect Immun 84, 2861–2870.

Kano, S.I., Hodgkinson, C.A., Jones-Brando, L., Eastwood, S., Ishizuka, K., Niwa, M., Choi, E.Y., Chang, D.J., Chen, Y., Velivela, S.D., et al. (2018). Host-parasite interaction associated with major mental illness. Mol Psychiatry.

Kawaguchi, Y. (1993). Groupings of nonpyramidal and pyramidal cells with specific physiological and morphological characteristics in rat frontal cortex. J Neurophysiol 69, 416–431.

Kawaguchi, Y., and Kubota, Y. (1997). GABAergic cell subtypes and their synaptic connections in rat frontal cortex. Cereb Cortex 7, 476–486.

Kawaguchi, Y., and Kubota, Y. (1998). Neurochemical features and synaptic connections of large physiologically-identified GABAergic cells in the rat frontal cortex. Neuroscience 85, 677–701.

Kettenmann, H., Kirchhoff, F., and Verkhratsky, A. (2013). Microglia: new roles for the synaptic stripper. Neuron 77, 10–18.

Koeniger, T., and Kuerten, S. (2017). Splitting the “Unsplittable”: Dissecting Resident and Infiltrating Macrophages in Experimental Autoimmune Encephalomyelitis. Int J Mol Sci 18.

Krasemann, S., Madore, C., Cialic, R., Baufeld, C., Calcagno, N., El Fatimy, R., Beckers, L., O’Loughlin, E., Xu, Y., Fanek, Z., et al. (2017). The TREM2-APOE Pathway Drives the Transcriptional Phenotype of Dysfunctional Microglia in Neurodegenerative Diseases. Immunity 47, 566–581 e569.

Kreutzfeldt, M., Bergthaler, A., Fernandez, M., Bruck, W., Steinbach, K., Vorm, M., Coras, R., Blumcke, I., Bonilla, W.V., Fleige, A., et al. (2013). Neuroprotective intervention by interferon-gamma blockade prevents CD8+ T cell-mediated dendrite and synapse loss. J Exp Med 210, 2087–2103.

Krook-Magnuson, E., Armstrong, C., Oijala, M., and Soltesz, I. (2013). On-demand optogenetic control of spontaneous seizures in temporal lobe epilepsy. Nat Commun 4, 1376.

Kuhlman, S.J., Olivas, N.D., Tring, E., Ikrar, T., Xu, X., and Trachtenberg, J.T. (2013). A disinhibitory microcircuit initiates critical-period plasticity in the visual cortex. Nature 501, 543–546.

Lambert, H., Hitziger, N., Dellacasa, I., Svensson, M., and Barragan, A. (2006). Induction of dendritic cell migration upon Toxoplasma gondii infection potentiates parasite dissemination. Cell Microbiol 8, 1611–1623.

Lang, D., Schott, B.H., van Ham, M., Morton, L., Kulikovskaja, L., Herrera-Molina, R., Pielot, R., Klawonn, F., Montag, D., Jansch, L., et al. (2018). Chronic Toxoplasma infection is associated with distinct alterations in the synaptic protein composition. J Neuroinflammation 15, 216.

Lewis, D.A., Fish, K.N., Arion, D., and Gonzalez-Burgos, G. (2011). Perisomatic inhibition and cortical circuit dysfunction in schizophrenia. Curr Opin Neurobiol 21, 866–872.

Li, Y., Severance, E.G., Viscidi, R.P., Yolken, R.H., and Xiao, J. (2019). Persistent Toxoplasma Infection of the Brain Induced Neurodegeneration Associated with Activation of Complement and Microglia. Infect Immun 87.

Marin, O. (2012). Interneuron dysfunction in psychiatric disorders. Nat Rev Neurosci 13, 107–120.

Marqueze, B., Boudier, J.A., Mizuta, M., Inagaki, N., Seino, S., and Seagar, M. (1995). Cellular localization of synaptotagmin I, II, and III mRNAs in the central nervous system and pituitary and adrenal glands of the rat. J Neurosci 15, 4906–4917.

Martin, H.L., Alsaady, I., Howell, G., Prandovszky, E., Peers, C., Robinson, P., and McConkey, G.A. (2015). Effect of parasitic infection on dopamine biosynthesis in dopaminergic cells. Neuroscience 306, 50–62.

McConkey, G.A., Martin, H.L., Bristow, G.C., and Webster, J.P. (2013). Toxoplasma gondii infection and behaviour - location, location, location? J Exp Biol 216, 113–119.

Mitra, R., Sapolsky, R.M., and Vyas, A. (2013). Toxoplasma gondii infection induces dendritic retraction in basolateral amygdala accompanied by reduced corticosterone secretion. Dis Model Mech 6, 516–520.

Monavarfeshani, A., Stanton, G., Van Name, J., Su, K., Mills, W.A., 3rd, Swilling, K., Kerr, A., Huebschman, N.A., Su, J., and Fox, M.A. (2018). LRRTM1 underlies synaptic convergence in visual thalamus. Elife 7.

Montoya, J.G., and Liesenfeld, O. (2004). Toxoplasmosis. Lancet 363, 1965–1976.

Mukherjee, A., Carvalho, F., Eliez, S., and Caroni, P. (2019). Long-Lasting Rescue of Network and Cognitive Dysfunction in a Genetic Schizophrenia Model. Cell 178, 1387–1402 e1314.

Neumann, J., Gunzer, M., Gutzeit, H.O., Ullrich, O., Reymann, K.G., and Dinkel, K. (2006). Microglia provide neuroprotection after ischemia. FASEB J 20, 714–716.

Ngoungou, E.B., Bhalla, D., Nzoghe, A., Darde, M.L., and Preux, P.M. (2015). Toxoplasmosis and epilepsy--systematic review and meta analysis. PLoS Negl Trop Dis 9, e0003525.

Palmer, B.S. (2007). Meta-analysis of three case controlled studies and an ecological study into the link between cryptogenic epilepsy and chronic toxoplasmosis infection. Seizure 16, 657–663.

Paolicelli, R.C., Jawaid, A., Henstridge, C.M., Valeri, A., Merlini, M., Robinson, J.L., Lee, E.B., Rose, J., Appel, S., Lee, V.M., et al. (2017). TDP-43 Depletion in Microglia Promotes Amyloid Clearance but Also Induces Synapse Loss. Neuron 95, 297–308 e296.

Pappas, G., Roussos, N., and Falagas, M.E. (2009). Toxoplasmosis snapshots: global status of Toxoplasma gondii seroprevalence and implications for pregnancy and congenital toxoplasmosis. Int J Parasitol 39, 1385–1394.

Parlog, A., Schluter, D., and Dunay, I.R. (2015). Toxoplasma gondii-induced neuronal alterations. Parasite Immunol 37, 159–170.

Paz, J.T., and Huguenard, J.R. (2015). Microcircuits and their interactions in epilepsy: is the focus out of focus? Nat Neurosci 18, 351–359.

Perry, V.H., and O’Connor, V. (2010). The role of microglia in synaptic stripping and synaptic degeneration: a revised perspective. ASN Neuro 2, e00047.

Peterson, J.W., Bo, L., Mork, S., Chang, A., and Trapp, B.D. (2001). Transected neurites, apoptotic neurons, and reduced inflammation in cortical multiple sclerosis lesions. Ann Neurol 50, 389–400.

Pfeffer, C.K., Xue, M., He, M., Huang, Z.J., and Scanziani, M. (2013). Inhibition of inhibition in visual cortex: the logic of connections between molecularly distinct interneurons. Nat Neurosci 16, 1068–1076.

Pi, H.J., Hangya, B., Kvitsiani, D., Sanders, J.I., Huang, Z.J., and Kepecs, A. (2013). Cortical interneurons that specialize in disinhibitory control. Nature 503, 521–524.

Poirotte, C., Kappeler, P.M., Ngoubangoye, B., Bourgeois, S., Moussodji, M., and Charpentier, M.J. (2016). Morbid attraction to leopard urine in Toxoplasma-infected chimpanzees. Curr Biol 26, R98–99.

Prandovszky, E., Gaskell, E., Martin, H., Dubey, J.P., Webster, J.P., and McConkey, G.A. (2011). The neurotropic parasite Toxoplasma gondii increases dopamine metabolism. PLoS One 6, e23866.

Savage, J.C., Picard, K., Gonzalez-Ibanez, F., and Tremblay, M.E. (2018). A Brief History of Microglial Ultrastructure: Distinctive Features, Phenotypes, and Functions Discovered Over the Past 60 Years by Electron Microscopy. Front Immunol 9, 803.

Schafer, D.P., Lehrman, E.K., Kautzman, A.G., Koyama, R., Mardinly, A.R., Yamasaki, R., Ransohoff, R.M., Greenberg, M.E., Barres, B.A., and Stevens, B. (2012). Microglia sculpt postnatal neural circuits in an activity and complement-dependent manner. Neuron 74, 691–705.

Schwaller, B., Tetko, I.V., Tandon, P., Silveira, D.C., Vreugdenhil, M., Henzi, T., Potier, M.C., Celio, M.R., and Villa, A.E. (2004). Parvalbumin deficiency affects network properties resulting in increased susceptibility to epileptic seizures. Mol Cell Neurosci 25, 650–663.

Sekar, A., Bialas, A.R., de Rivera, H., Davis, A., Hammond, T.R., Kamitaki, N., Tooley, K., Presumey, J., Baum, M., Van Doren, V., et al. (2016). Schizophrenia risk from complex variation of complement component 4. Nature 530, 177–183.

Sgado, P., Dunleavy, M., Genovesi, S., Provenzano, G., and Bozzi, Y. (2011). The role of GABAergic system in neurodevelopmental disorders: a focus on autism and epilepsy. Int J Physiol Pathophysiol Pharmacol 3, 223–235.

Shi, Y., Yamada, K., Liddelow, S.A., Smith, S.T., Zhao, L., Luo, W., Tsai, R.M., Spina, S., Grinberg, L.T., Rojas, J.C., et al. (2017). ApoE4 markedly exacerbates tau-mediated neurodegeneration in a mouse model of tauopathy. Nature 549, 523–527.

Sims, T.A., Hay, J., and Talbot, I.C. (1989). An electron microscope and immunohistochemical study of the intracellular location of Toxoplasma tissue cysts within the brains of mice with congenital toxoplasmosis. Br J Exp Pathol 70, 317–325.

Siskova, Z., Page, A., O’Connor, V., and Perry, V.H. (2009). Degenerating synaptic boutons in prion disease: microglia activation without synaptic stripping. Am J Pathol 175, 1610–1621.

Skallova, A., Kodym, P., Frynta, D., and Flegr, J. (2006). The role of dopamine in Toxoplasma-induced behavioural alterations in mice: an ethological and ethopharmacological study. Parasitology 133, 525–535.

Stevens, B., Allen, N.J., Vazquez, L.E., Howell, G.R., Christopherson, K.S., Nouri, N., Micheva, K.D., Mehalow, A.K., Huberman, A.D., Stafford, B., et al. (2007). The classical complement cascade mediates CNS synapse elimination. Cell 131, 1164–1178.

Stock, A.K., Dajkic, D., Kohling, H.L., von Heinegg, E.H., Fiedler, M., and Beste, C. (2017). Humans with latent toxoplasmosis display altered reward modulation of cognitive control. Sci Rep 7, 10170.

Stock, A.K., Heintschel von Heinegg, E., Kohling, H.L., and Beste, C. (2014). Latent Toxoplasma gondii infection leads to improved action control. Brain Behav Immun 37, 103–108.

Su, J., Chen, J., Lippold, K., Monavarfeshani, A., Carrillo, G.L., Jenkins, R., and Fox, M.A. (2016). Collagen-derived matricryptins promote inhibitory nerve terminal formation in the developing neocortex. J Cell Biol 212, 721–736.

Su, J., Gorse, K., Ramirez, F., and Fox, M.A. (2010). Collagen XIX is expressed by interneurons and contributes to the formation of hippocampal synapses. J Comp Neurol 518, 229–253.

Tang, Y., Stryker, M.P., Alvarez-Buylla, A., and Espinosa, J.S. (2014). Cortical plasticity induced by transplantation of embryonic somatostatin or parvalbumin interneurons. Proc Natl Acad Sci U S A 111, 18339–18344.

Torres, L., Robinson, S.A., Kim, D.G., Yan, A., Cleland, T.A., and Bynoe, M.S. (2018). Toxoplasma gondii alters NMDAR signaling and induces signs of Alzheimer’s disease in wild-type, C57BL/6 mice. J Neuroinflammation 15, 57.

Trapp, B.D., Wujek, J.R., Criste, G.A., Jalabi, W., Yin, X., Kidd, G.J., Stohlman, S., and Ransohoff, R. (2007). Evidence for synaptic stripping by cortical microglia. Glia 55, 360–368.

Tremblay, M.E., Lowery, R.L., and Majewska, A.K. (2010). Microglial interactions with synapses are modulated by visual experience. PLoS Biol 8, e1000527.

Tremblay, R., Lee, S., and Rudy, B. (2016). GABAergic Interneurons in the Neocortex: From Cellular Properties to Circuits. Neuron 91, 260–292.

Tukker, J.J., Fuentealba, P., Hartwich, K., Somogyi, P., and Klausberger, T. (2007). Cell type-specific tuning of hippocampal interneuron firing during gamma oscillations in vivo. J Neurosci 27, 8184–8189.

Vasek, M.J., Garber, C., Dorsey, D., Durrant, D.M., Bollman, B., Soung, A., Yu, J., Perez-Torres, C., Frouin, A., Wilton, D.K., et al. (2016). A complement-microglial axis drives synapse loss during virus-induced memory impairment. Nature 534, 538–543.

Vyas, A., Kim, S.K., Giacomini, N., Boothroyd, J.C., and Sapolsky, R.M. (2007). Behavioral changes induced by Toxoplasma infection of rodents are highly specific to aversion of cat odors. Proc Natl Acad Sci U S A 104, 6442–6447.

Wang, Z.T., Harmon, S., O’Malley, K.L., and Sibley, L.D. (2015). Reassessment of the role of aromatic amino acid hydroxylases and the effect of infection by Toxoplasma gondii on host dopamine. Infect Immun 83, 1039–1047.

Wang, Z.T., Verma, S.K., Dubey, J.P., and Sibley, L.D. (2017). The aromatic amino acid hydroxylase genes AAH1 and AAH2 in Toxoplasma gondii contribute to transmission in the cat. PLoS Pathog 13, e1006272.

Wang, A.W., Avramopoulos, D., Lori, A., Mulle, J., Conneely, K., Powers, A., Duncan, E., Almli, L., Massa, N., McGrath, J., et al. (2019). Genome-wide association study in two populations to determine genetic variants associated with Toxoplasma gondii infection and relationship to schizophrenia risk. Prog Neuropsychopharmacol Biol Psychiatry 92, 133–147.

Weinhard, L., di Bartolomei, G., Bolasco, G., Machado, P., Schieber, N.L., Neniskyte, U., Exiga, M., Vadisiute, A., Raggioli, A., Schertel, A., et al. (2018). Microglia remodel synapses by presynaptic trogocytosis and spine head filopodia induction. Nat Commun 9, 1228.

Wohlfert, E.A., Blader, I.J., and Wilson, E.H. (2017). Brains and Brawn: Toxoplasma Infections of the Central Nervous System and Skeletal Muscle. Trends Parasitol 33, 519–531.

Wohr, M., Orduz, D., Gregory, P., Moreno, H., Khan, U., Vorckel, K.J., Wolfer, D.P., Welzl, H., Gall, D., Schiffmann, S.N., et al. (2015). Lack of parvalbumin in mice leads to behavioral deficits relevant to all human autism core symptoms and related neural morphofunctional abnormalities. Transl Psychiatry 5, e525.

Xiao, J., Prandovszky, E., Kannan, G., Pletnikov, M.V., Dickerson, F., Severance, E.G., and Yolken, R.H. (2018). Toxoplasma gondii: Biological Parameters of the Connection to Schizophrenia. Schizophr Bull 44, 983–992.

Yamasaki, R., Lu, H., Butovsky, O., Ohno, N., Rietsch, A.M., Cialic, R., Wu, P.M., Doykan, C.E., Lin, J., Cotleur, A.C., et al. (2014). Differential roles of microglia and monocytes in the inflamed central nervous system. J Exp Med 211, 1533–1549.

