## Supplementary figures and images for "*Toxoplasma* induces stripping of perisomatic inhibitory synapses"

### Supplemental Figure 1

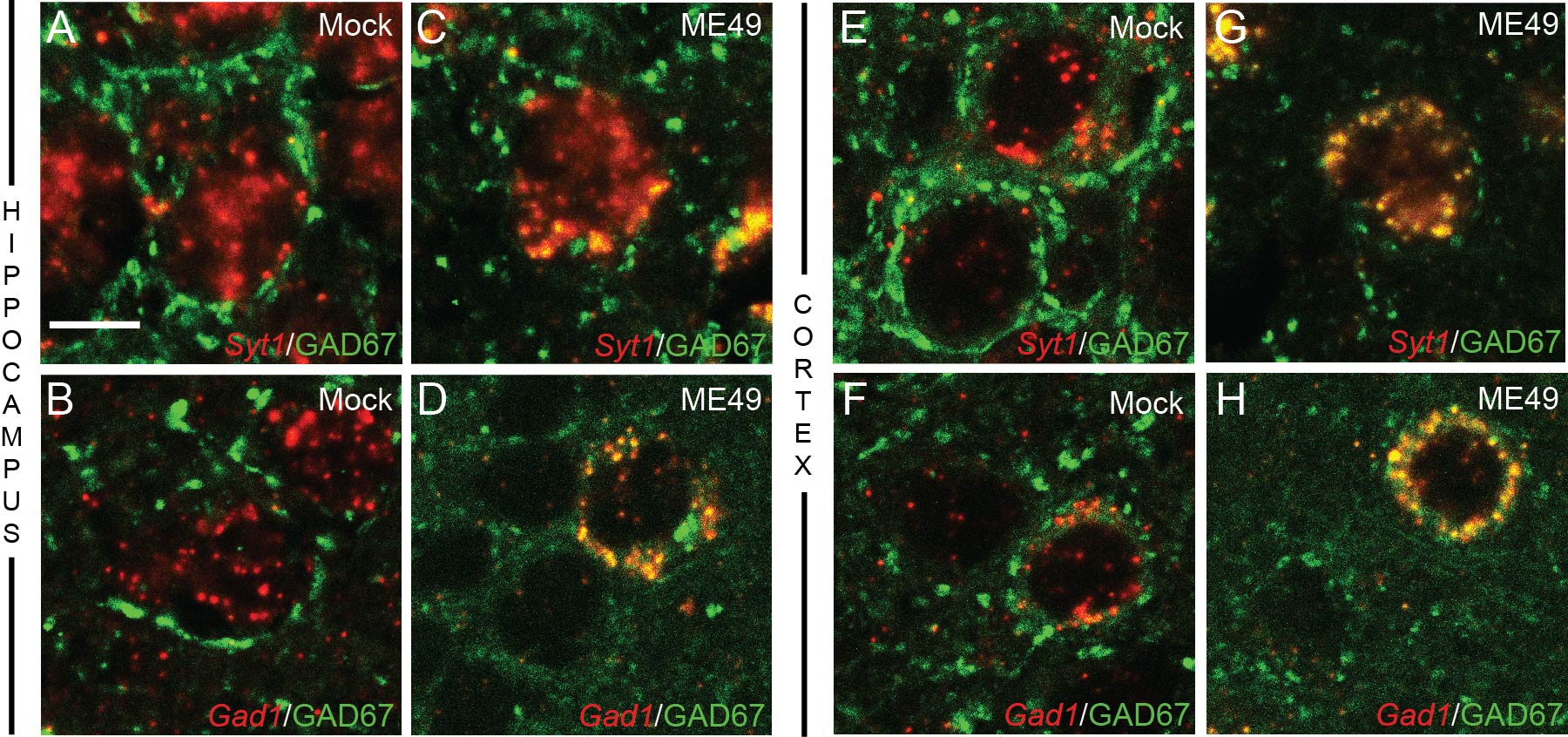

### Supplemental Figure 2

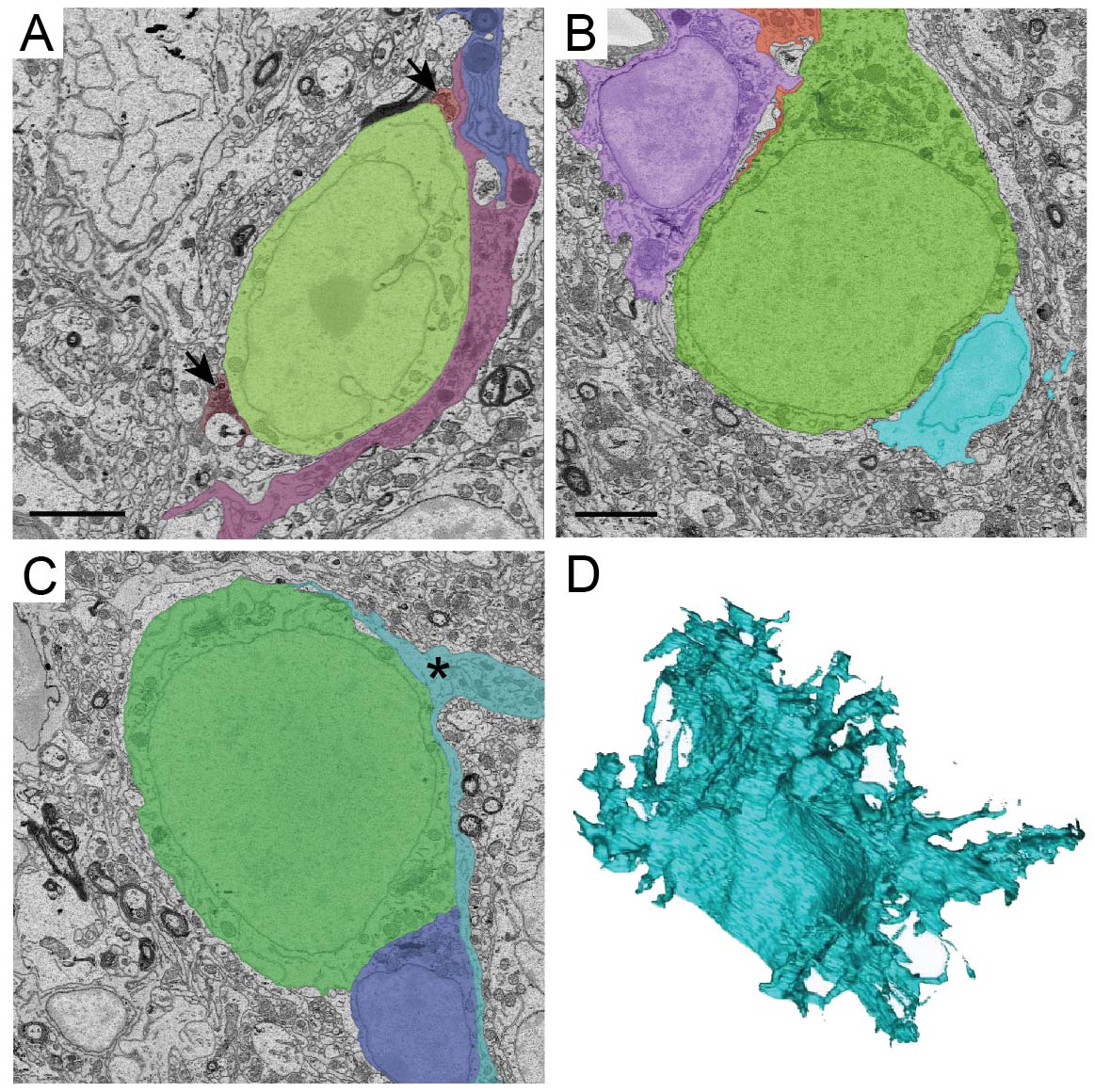

### Supplemental Figure 3

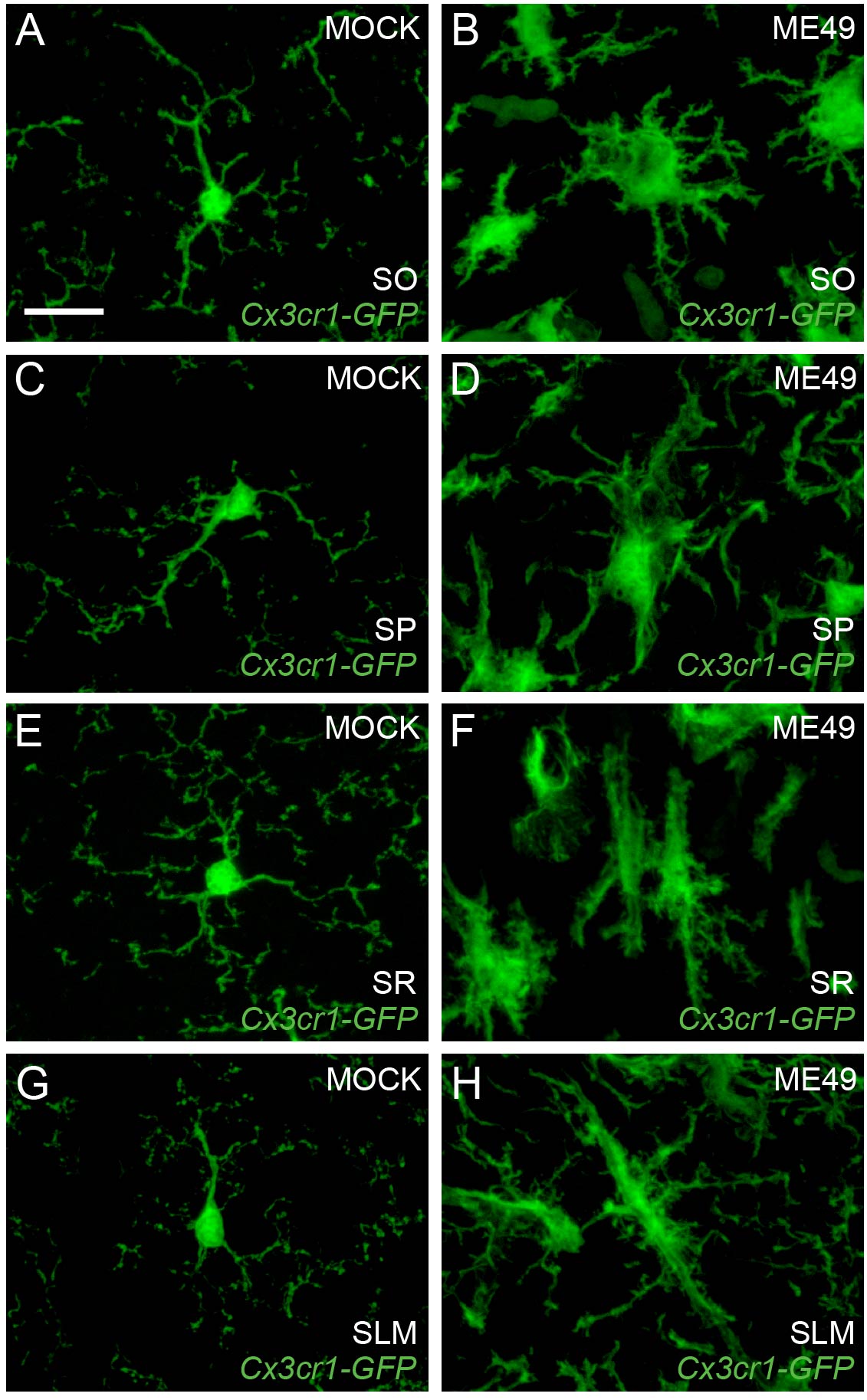
